# Human fallopian tube-on-a-chip for preclinical testing of non-hormonal contraceptives with living human sperm

**DOI:** 10.64898/2026.01.22.700844

**Authors:** Anna Stejskalová, Karina Calderon, McKenzie Collins, Jessica F Feitor, Debraj Ghose, Shiyan Tang, Ola Gutzeit, Nikil Badey, Aakanksha Gulati, Maria V Lopez, David B Chou, John C Petrozza, Roberto Plebani, Abidemi Junaid, Bogdan Budnik, Donald E Ingber

**Author notes:** These authors contributed equally as second authors (Karina Calderon, McKenzie Collins). Senior author (Donald E Ingber).

## Abstract

The fallopian tube serves as a sperm reservoir, and it is the site where the oocytes become fertilized. Here, we describe development of an organ-on-a-chip microfluidic model of the fallopian tube (FT Chip) lined by primary human epithelial cells and stromal fibroblasts derived from the FT ampulla. Abundant tissue folds lined by hormone-responsive, epithelial cells resembling those seen *in vivo* formed on-chip, but not in epithelial organoids cultured in gel cultures. Comparative time-resolved analysis of human sperm versus oocyte-sized microparticles introduced into the epithelial channel in the presence of estradiol revealed that sperm movement was significantly reduced, while the oocyte-sized particles increased, relative to movements in acellular chips. When the non-hormonal contraceptive TDI-11861 was administered to the chip, dose-dependent inhibition of human sperm motility was detected. Thus, this FT Chip may offer a human preclinical tool to study FT physiology and assess the efficacy and mechanism of action of contraceptives.

## INTRODUCTION

The fallopian tube (FT) plays a central role in human reproduction, providing the site of fertilization and supporting early embryo transport^1^. In lower mammalian species, the ampulla region of the FT also serves as a sperm reservoir^2^, where ciliated epithelial surfaces transiently bind spermatozoa and sustain them until ovulation, when cumulus-oocyte-complex (C-O-C) cues trigger sperm hyperactivation and fertilization^3,4^. In contrast, the cellular and functional dynamics of the human FT remain largely unexplored, including whether a comparable sperm reservoir exists and how the FT’s remarkable hormone responsiveness influences fertilization efficiency^5^. These questions remain unanswered because the FT is one of the least accessible human tissues, precluding live imaging, fluid sampling, or direct live observation. This limitation has impeded efforts to develop precision strategies for preventing FT disease, ranging from infertility, ectopic pregnancy to malignancy. In recent years, precision female-controlled contraceptives have become recognized as a key primary prevention strategy to decrease maternal mortality in developing countries^6^, yet the efficacy of these strategies, many of which are effective primarily in the FT^7^, are challenging to evaluate. Thus, this basic science and pre-clinical bottleneck, which has resulted in a lack of suitable contraceptive options, has significant global health implications.

The FT consists of four main parts, the isthmus, ampulla, infundibulum and fimbrae^8^ that have distinct functions and are derived from distinct epithelial lineages^9^. Morphologically, the FT epithelium is organized into abundant branched folds, consisting primarily of secretory and ciliated lineages. with proliferative cells being distributed along the folds^10^. Cohort and single-cell RNA sequencing studies have revealed that the FT epithelium is highly dynamic, assuming distinct molecular states and architecture throughout the menstrual cycle and following the menopause^11^. However, the functional implications of these profound FT transitions on human reproduction and disease remain to be elucidated.

Stem cell-derived models offer a promising strategy to overcome the challenges related to the inaccessibility of the human FT for direct observation and manipulation in a controlled manner. Although primary human FT stem cell-based organoids replicate key epithelial lineages^10^, they fail to reproduce the complex folded architecture that guides sperm translocation *in vivo*^12^, and their closed geometry^13^ precludes co-culture with sperm or oocytes. Transwell culture^14^ does not enable continuous hormonal perfusion and lacks the optical transparency required for live imaging; it is also difficult to analyze directional sperm movement. Recent development of microfluidic platforms, including organ-on-a-chip (Organ Chip) technologies^15^ using cells from non-human oviducts^16^ have begun to recapitulate aspects of reproductive tract physiology, yet none capture the organ-specific tissue architecture, cellular composition, and hormone-regulated microenvironment of the human FT.

To address these limitations, we developed a primary human fallopian tube-on-a-chip microfluidic culture model (FT Chip) lined by primary human epithelial cells and stromal fibroblasts derived from the FT ampulla that reconstitutes the epithelial folds, tubal architecture, and dynamic ciliary function of the native tissue under continuous physiological hormone flow. This platform enables direct, long-term co-culture with human spermatozoa and provides a physiologically relevant environment for assessing the efficacy and mechanisms of action of candidate non-hormonal contraceptives within a human organ-specific reproductive context.

## RESULTS

### Fallopian Tube Chips replicate the folded epithelial architecture of the native ampulla

To develop a human FT Chip, we combined advances in microfluidic Organ Chip and adult stem cell-derived epithelial organoid technologies. To ensure relevance for fertility studies, we specifically re-created the ampulla part of the FT that is the key region of the organ that serves as the meeting point for the sperm and oocyte during human fertilization. The FT was donated by hysterectomy patients and processed by an experienced pathologist, and epithelial stem cells and matched fibroblasts were isolated from the same region. The epithelial cells were expanded in conventional planar culture dishes using a previously described protocol^10^ and then either grown as organoids within Matrigel cultures^10^ and used to create FT Chips.

The FT Chips were created by co-culturing the primary FT epithelial cells on top of a porous ECM-coated membrane in the apical channel of a commercially available two-channel Organ Chip and interfacing them with matched fibroblasts from the same donor, which were cultured on the bottom of the same membrane in a parallel channel (**Figure 1A**). Both cell layers were cultured in growth medium under dynamic flow (30 μl/hr) for 4 days to confluence. Epithelial differentiation was then induced by perfusing Human Tubal Fluid (HTF) medium through the apical channel for 4 hours (40 µl/hr) each day while nutrient-rich differentiation medium (Gynecult) was continually perfused (45µl/hr) through the basal channel in the presence of the steroid hormone estradiol (E2). HTF was chosen because it is used clinically to support sperm function during human *In vitro* fertilization (IVF) procedures as it contains components found in native FT fluid, including essential electrolytes, albumin and energy sources (e.g., glucose)^17^. Several basal differentiation media were evaluated (**Supplementary Figure S1**), with the commercially available Gynecult medium resulting in the most uniform phenotype across the full 2 cm length of the epithelial channel in the FT Chip. FT Chips differentiated under these conditions formed abundant folds typical for this tissue *in vivo* (**Figure 1B**).

**Figure 1.**
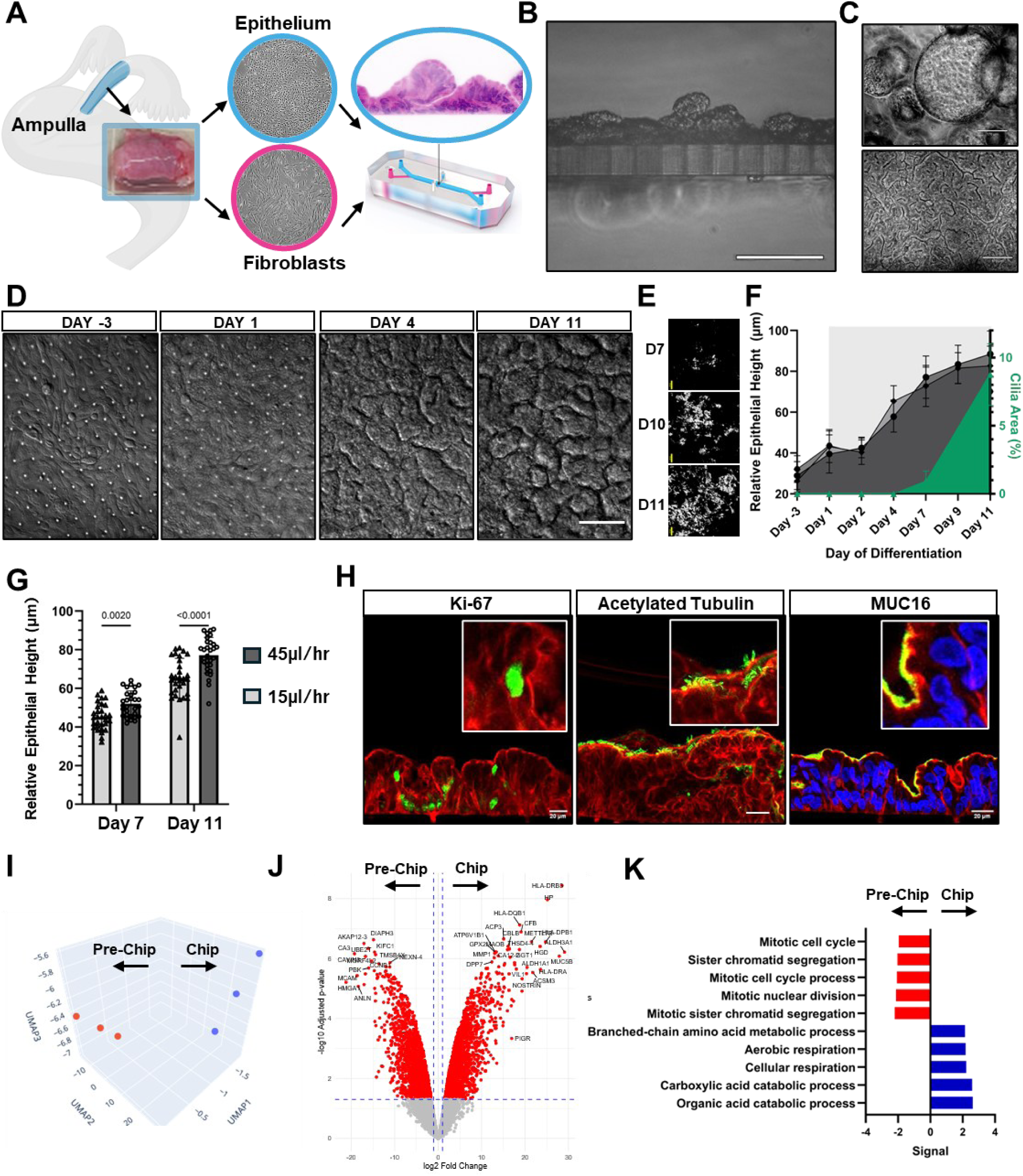
Fallopian Tube Chips replicate the folded epithelial architecture of the native ampulla. (A) A schematic of the Ampulla region of the Fallopian tube and the Fallopian Tube Organ Chip. Created in BioRender. Chip photograph source www.emulatebio.com (B) Microscopic image depicting a cross-section of the Fallopian Tube Chip (bar, 120 µm). (C) Phase contrast microscopic images of fallopian tube epithelial organoids (top) and a FT Chip cultured in parallel using the same media on day 16 of culture (bar, 170 µm). (D) Representative DIC microscopic images of FT Chips differentiated under E2 perfusion and fixed on day 2, 6, 9 or 16 of culture (bar, 100 µm). (E) Representative images of segmented beating cilia on days 7, 10 and 11 of differentiation (bar, 20 µm). See Video S1. (F) Graph showing changes in the height of the epithelial layer over time (left Y-axis) and in the area covered by ciliated epithelial cells (right Y-axis, green). The individual height repeats visualized as mean±SD represent three biological donor replicates, each calculated from an N=4 chips, 10 sampling locations per chip. Cilia levels were determined using one donor. (G) A bar graph showing epithelial heights measured in FT Chips perfused basally at low (15μL/h; light grey) or high (45 μL/h; dark grey) flow rates (N = 3 chips; one biological replicate, 10 sampling locations per chip; p values are determined using two-way ANOVA). (H) Immunofluorescence microscopic images of FT Chip cross-sections depicting the F-actin cytoskeleton (red) stained with fluorescent phalloidin. The left image shows nuclei (green, Hoechst), middle image cilia (green, acetylated tubulin) and the right MUC16 (green);(bar, 20 µm). (I) UMAP 3D plot of the bulk proteomic data obtained from FT epithelial cells cultured in 2D (Pre-Chip, Red) versus same epithelial cells following their expansion on-chip in the absence of added steroid hormones (Chip, blue) (N=3 Chips, N=1 biological donor). (J) Volcano plot depicting differentially expressed proteins in epithelia cultured in 2D (pre-chip, N=3 technical mass spectrometry aliquot repeats, N=1 biological donor) and harvested from chips on day 16 of culture (N=3 chips, N=1 biological donor) (K) Graph depicting top 5 Biological Processes upregulated in the FT Chips cultured without steroid hormone compared to cells cultured in 2D generated from bulk proteomics analysis of differentially expressed proteins with adjusted p value<0.01.

Interestingly, the folded phenotype appears to uniquely arise in planar polarized cultures, as when we analyzed organoids created the same human FT ampulla-derived epithelial cells that were cultured within Matrigel under static conditions in the same medium, we only observed cystic structures that failed to form folds even after 16 days of culture (**Figure 1C**), which aligns with results of published FT organoid studies^18^.

Time-resolved analysis of the FT Chips lined by cells from three different human donors using differential interference contrast (DIC) imaging revealed that the differentiation timecourse in the presence of E2 is highly conserved (**Figure 1D**). In particular, the epithelium started to form apically protruding folds between 2 and 4 days after being switched to the differentiation conditions, and these folds continued to increase in height up until day 7 when the height increase slowed. Importantly, populations with functional beating cilia also appeared and became increasingly abundant between days 7 and 11 of differentiation as evidenced from segmented video microscopy data (**Figure 1E, F, Video S1**).

When we compared the use of high versus low basal medium flow rates (45 versus 15 µL/hr, respectively), we found that faster flow promoted formation of taller and wider epithelial folds (**Figure 1G**).

Immunohistochemical analysis confirmed that the epithelium lining the FT Chips developed a folded morphology (**Figure 1H**) closely resembling the native differentiated architecture of the human FT ampulla. We also confirmed that the FT chip epithelia remain proliferative, as evidenced by Ki-67 nuclear localization even on day 16 of culture. Moreover, the diffuse localization of Ki-67 positive cells along the epithelial folds on-chip parallels the diffuse location of proliferative lineages in the native FT^10^. The FT Chips also develop abundant cilia (**Figure 1H**) and contain abundant MUC16^+^ (CA125) lineages.

Uniform Manifold Approximation and Projection (UMAP) analysis of the proteomic data revealed that the changes in the FT epithelium observed on-chip were not only morphological, but also biochemical (**Figure 1I**), resulting in a large number of differentially expressed proteins compared to 2D culture (**Figure 1J**). Differential bulk epithelial proteome and gene ontology analysis of samples from the FT epithelial cells from the same donor grown under differentiation conditions on-chip versus in conventional two-dimensional (2D) culture dishes revealed a marked shift in cell metabolism, transitioning from a cancer-resembling proliferative signature in 2D (gene ontology terms: ‘*Mitotic cell cycle’*, ‘*Sister chromatid segregation’*) to being enriched in processes corresponding to the Biological Processes ‘*Aerobic respiration’*, and ‘*Carboxylic acid and organic acid catabolic processes*’ in the FT Chips (**Figure 1K**). Many of the top fifteen upregulated proteins are considered stem cell or cancer stem cell markers, including ALDH3A1 and ALDH1A1^19^. One notable protein that was upregulated in the differentiated epithelium on-chip included epididymis-specific alpha-mannosidase (MAN2B2-2), an enzyme that is involved in the maturation of spermatozoa in the male reproductive tract by deglycosylating the sperm head^20^. The epithelium in the FT Chip also upregulated the levels of the sodium-bicarbonate cotransporter SLC4A4 that is involved in basolateral bicarbonate uptake; this is important because apical bicarbonate levels are essential for increasing local pH and inducing sperm capacitation. The epithelium grown in 2D culture, on the other hand, was marked by higher expression of MUC18 (MCAM), a cell surface glycoprotein with known functions during melanoma tumor spreading^21^ (**Supplementary Figure 2**).

#### Fallopian Tube Chips are hormone-responsive

The native FT adopts distinct biological functions throughout the menstrual cycle (**Figure 2A**). To evaluate whether the FT Chip recapitulates hormone responsiveness seen *in vivo*^22^, the FT chips were differentiated in the absence of hormones (no hormones, NH), in the presence of 2nM Estradiol (E2), or the presence of 0.1 M E2 and 1 µM Progesterone (E2+P4) to mimic the menstrual, follicular and luteal phases, respectively. The E2 group had significantly different morphology from NH controls and the E2+P4 group (**Figure 2B,C**) with E2 exposure alone significantly increasing both the relative height and width of the folds (**Figure 2D, E**). The folds that developed under NH covered only a minimal area compared to the E2 group. In addition, while the FT Chip morphology was comparable across three human donors when differentiated in the presence of E2, the response was highly variable across donors in the absence of hormones (**Supplementary Figure 3A, B**).

**Figure 2.**
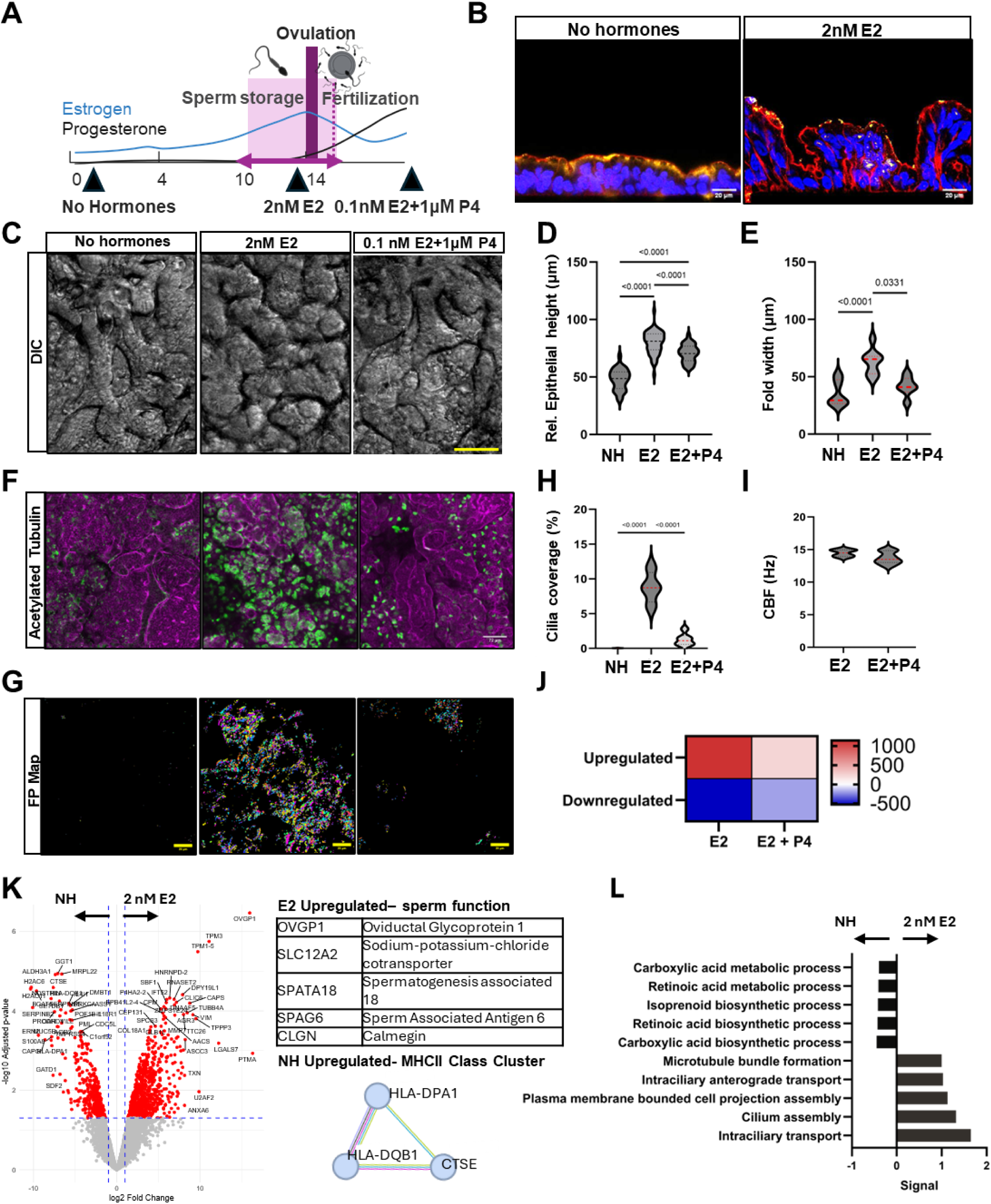
Fallopian Tube Chips are hormone responsive. (A) Diagram depicting the fertile window and shifts in circulating hormone levels related to the menstrual cycle. Times when the E2 level are highest correspond to the “Fertile Window” (light pink) portion of the follicular phase when likelihood of fertilization is highest. Created in BioRender. (B) Immunofluorescence microscopic images of cross-sections of the human FT Chip perfused basally with medium containing differentiation medium alone (no hormones) with E2 (2 nM) on day 16 stained for the F-actin (red), PAX8 (yellow) and nuclei (blue) (bar, 20 µm). (C) DIC microscopic image of FT epithelia on-chip perfused with medium with or without E2 hormone (bar, 100µm). (D) Violin plots depicting the relative height of the epithelial layer on FT Chips when cultured for 16 days with no hormones (NH), E2 alone, or E2 + P4 (N = 2 donors, N=3 chips per condition and donor, 10 sampling points per chip, p values were determined using a one way-ANOVA). (E) Violin plot depicting the width of epithelial folds on day 16 (N=2 donors, 3-4 chips per condition and donor) (F) Reconstructions of immunofluorescence confocal microscopic images of FT Chips differentiated under different hormonal conditions and viewed from above on day 16 (F-actin, magenta; cilia, green; bar, 72 µm) (G) Images depict the frequency phase map showing representative relative cilia beating frequencies CBF (Hz) (H) Violin plots depicting the area covered by actively beating cilia (N=7, 4 and 6 chips per group. P values were determined using a one way-ANOVA) (J) Violin plot showing the median frequency of cilia beats for chips perfused with E2 or E2+P4 for 16 days (N = 4 and 6 chips respectively p values were determined using t-test). (J) Heatmap depicting a number of differentially expressed proteins between E2 perfused chips and controls and between E2+P4 perfused group and controls determined using differential protein expression analysis shown in Figure 2K (N=3 chips per condition, one biological donor) (K) Volcano plot of the cell epithelial proteome of FT Chips differentiated with E2 versus without steroid hormone (NH). The table lists upregulated proteins in the E2 group, and the String Network depicts a cluster identified with a MCL clustering algorithm (N=3 chip replicates from one biological donor). (L) A diverging bar chart showing the top five enriched biological processes in the proteomes of epithelial cells cultured in FT Chips cultured with no hormones (NH) versus E2 (normalized p value <0.1).

One of the most recognized roles of E2 in the FT is the induction of ciliogenesis^11,13,23^. Indeed, staining of the FT Chips with acetylated tubulin (**Figure 2F**) and live video imaging of actively beating cilia (**Figure 2G**) confirmed that perfusion with E2 significantly increases the number of ciliated cells compared with the NH and E2+P4 groups (**Figure 2H**). However, while the E2+P4 group had fewer cilia than the E2 group, the median ciliary beat frequency (CBF) did not significantly differ (**Figure 2I**). This finding contrasts with previous studies where progesterone decreased CBF in the mouse fallopian tube. This could be due to long-term exposure of the epithelium to dynamic flow of hormones from its basal surface on-chip compared to acute apical exposure used in past studies^24^ ^25^.

E2 also induced large and significant changes at the bulk epithelial cell proteome level compared to controls (**Figure 2J**) while treatment with E2+P4 resulted in relatively smaller effect compared to controls (**Figure 2J, Supplementary Figure 4A)** as identified using differential protein expression analysis. In agreement with prior *in vivo* and *in vitro* findings^16^, one of the proteins most upregulated in response to E2 was Oviductal Glycoprotein 1 (OVGP1), and other proteins potentially involved in specialized oviductal secretions (e.g., LGALS7, CAPS) were upregulated as well (**Figure 2K**). The top fifteen most differentially expressed proteins in the E2 group included cytoskeletal proteins, such as tropomyosins (TPM3, TPM1, TPM5), TPPP3, TUBB4A, VIM and TTC26 (**Supplementary Figure 4B**), which aligns with the significant increase in epithelial folding and ciliation observed in the presence of E2. Other differentially expressed proteins included molecular chaperones and stress response proteins, including DNAJA1, TXN, VBP1, ASCC3 and DPY19L1. In contrast, proteins more highly expressed in the NH FT Chips included major histocompatibility antigens (e.g., HLA-DRB3,HLA-DPA1) as well as CTSE and S100A8, which are involved in immune recognition and antigen presentation; MUC5B and ALDH3A1, involved in barrier and oxidative defenses; and CAPG and NOSTRIN that may be involved in regulating the cytoskeleton and motility. Thus, the absence of E2 appears to significantly alter the immune landscape within the FT, which aligns with recent single cell RNA-seq studies on native tissues showing that MHC-II transcripts are upregulated during the luteal and secretory phases^8^.

Overall, the presence of the key follicular stage hormone E2 appeared to upregulate proteins that support reproduction. For example, perfusion of the FT Chip with E2 resulted in significantly higher levels of proteins that may be relevant for optimal sperm function (**Figure 2K, Supplementary Figures 5 and 6**). These upregulated proteins include a sodium-potassium-chloride cotransporter (SLC12A2), spermatogenesis associated 18 (SPATA18) involved in mitochondrial quality control, sperm associated antigen 6 (SPAG6), a structural protein of the axoneme found in motile cilia or flagella, and Calmegin (CLGN), an ER-resident molecular chaperone for sperm glycoproteins that is essential for sperm fertility^26^.

Gene ontology analysis of biological processes (**Figure 2L**) conducted on the dataset shown in **Figure 2K** revealed that the most upregulated terms in the presence of E2 are related to cilia function, including ‘*Intraciliary transport’*, ‘*Cilium assembly’* and ‘*Membrane bounded cell projection assembly*’, which aligns with our immunohistochemical results (**Figure 2F)**. The absence of steroid hormones increased terms related to ‘*Carboxylic acid biosynthetic process*’ and ‘*Retinoic acid biosynthetic process’*. This is interesting as retinoic acid is essential for Müllerian duct development and epithelial homeostasis of the adult female reproductive tract^27^.

### Fallopian Tube Chips support different movements of sperm versus oocyte-sized particles

The ampulla serves as a site within the FT where the sperm fertilizes the oocyte. While fertilization can also happen *in vitro*, the role of FT epithelium in enabling this process *in vivo* and why fertilization always occurs in this specific anatomical location remains poorly understood. Currently, this has been attributed to regional variations in the biochemical species (e.g., ions^28^, proteins^29^) as well as physical conditions (e.g, spatial confinement^30^, sperm migration along fold walls and corners^31^, local viscosity^32^). To directly evaluate the contribution of the FT epithelium, we first compared movement of sperm head and oocyte sized fluorescent microparticles (5 and 100 µm diameter, respectively) when placed in the upper channel of epithelium-lined FT Chips differentiated in either the absence (No Hormones, NH) or presence of E2 (corresponding to the fertile window) versus acellular (empty) chips (**Figure 3A**).

**Figure 3.**
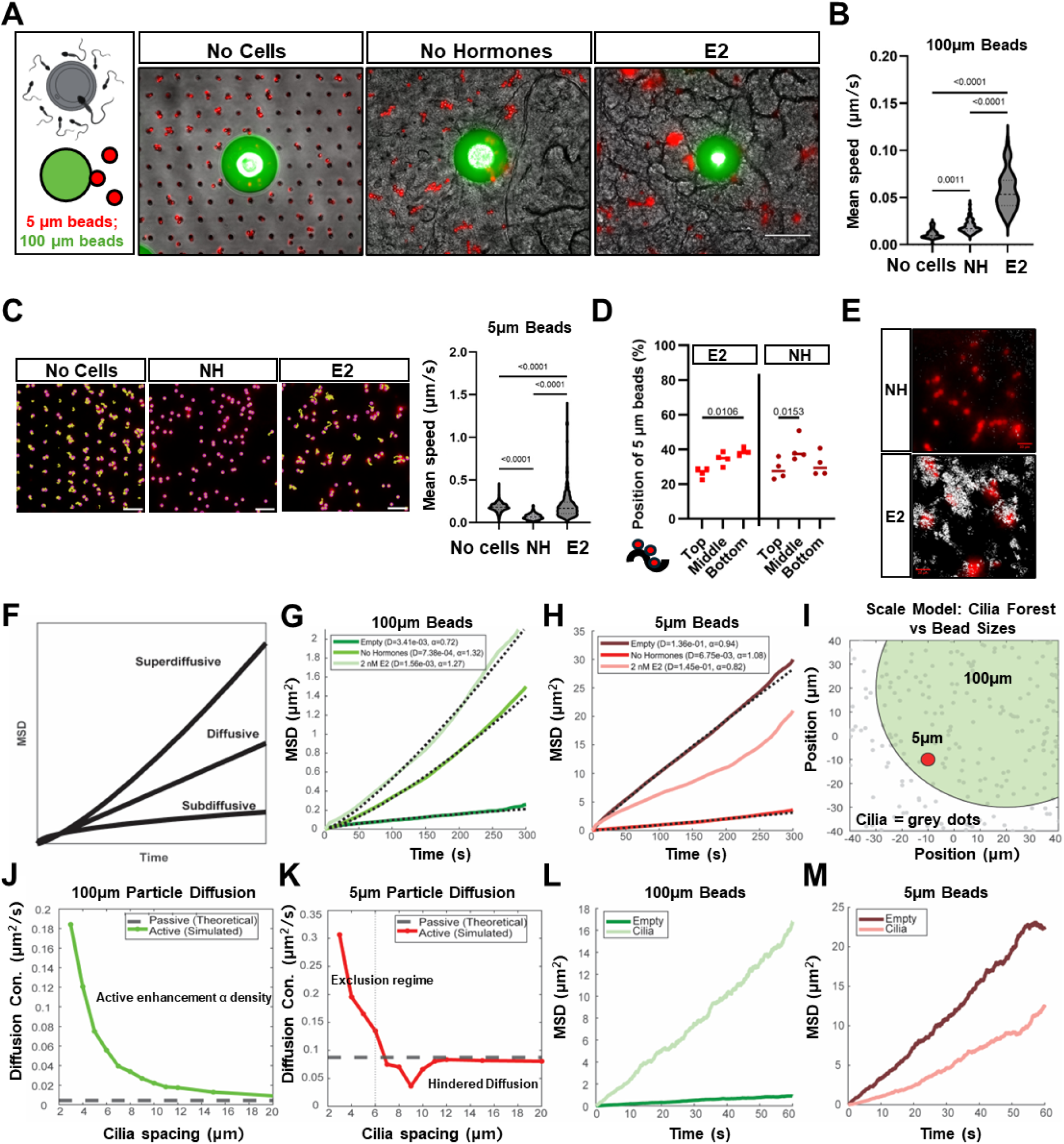
Follicular-Stage FT CHIP supports superdiffusive movement of oocyte-sized beads but subdiffusive motion of spermatozoa-sized particles. (A) A schematic figure of 100 and 5µm beads selected to mimic the oocyte (green) or spermatozoa head (red) (left). Created in BioRender. Representative microscopic images of 5 and 100 µm beads on an acellular chip (left) versus FT Chips lined by epithelium differentiated in the absence (middle) or presence (right) of E2. (B) Violin plot showing the parameter mean speed of 100 µm beads quantified on-chip under the conditions shown in A ( N=4, 5 and 5 chips, 66-73 beads were analyzed per group; p values determined by one way-ANOVA) (C) Computerized morphometric analysis of microscopic images showing representative movement tracks (yellow) of 5 µm beads (purple) measured on chips cultured without cells (Scale bar, 50µm), or with cells in the absence or presence of E2 (left) and violin plot showing quantification of mean speeds measured for these beads (right) (N= 4, 5 and 5 chips, 397-569 beads analyzed per group; p values determined by one way-ANOVA). See videos S2-S.4 (D) Graph visualizing the position of 5 µm beads along the fold height (z-axis) of the FT Chip visualized at three discrete levels (bottom, middle, top) cultured in the absence (•) or presence of E2 (▪). (Mean±SD; N= 4 chips; p values determined by one way-ANOVA) (E) Representative overlay images of 5 µm beads (red) and segmented beating cilia (white) in human FT Chips perfused without (top) or with E2 (bottom). Corresponding Videos S7-8. (F) Diagram depicting mean square displacement (MSD, Y axis) over time (X axis) for diffusive, superdiffusive and subdiffusive motion. (G) MSD plots (Y axis) depicting trajectories of 100µm beads from Figure 3B on empty (dark green), NH (medium green) and E2 FT (light green) chips over time (X axis). (H) MSD plots depicting trajectories of 5µm beads from Figure 3C on empty (dark red), NH (medium red) and E2 FT (light red) chips over time. In both G and H, the dotted line represents a fit to the equation msd = D*t^\alpha, D is a generalized diffusion prefactor rather than the physical diffusion coefficient. (I) A schematic depiction of simulation showing cilia (grey dots) at varying densities and beads of two different sizes (5µm, 100µm). (J) Predicted particle diffusion constant (Y axis) at different cilia spacing conditions (X axis) of 100µm particles under active propulsion (green) and passive (grey) condition. (K) Predicted particle diffusion constant (Y axis) at different cilia spacing conditions (X axis) of 5µm particles under active propulsion (red) and passive (grey) condition. (L) Simulated MSD over time of 100 µm beads in the presence (light green) or absence of cilia (dark green). (M) Simulated MSD over time of 5µm beads in the presence (light red) or absence of cilia (dark red).

Inorganic microparticles in fluid medium are subject to Brownian motion, whereby the diffusion coefficient (D) of a spherical particle under a constant temperature (T) is inversely proportional to particle radius (r) and fluid viscosity (η) (D=kB•T/6•π•η•r) (**Equation 1**). Thus, under otherwise equal conditions, diffusion should be significantly lower for the oocyte-sized 100 µm beads compared to the sperm-sized 5 µm particles. Indeed, single particle tracking revealed that the velocity of the oocyte-sized microbeads was lower than that of the smaller beads when introduced into the empty (acellular) Organ Chip devices in HTF medium (**Figure 3B,C**). The speed of the oocyte-sized microbeads was significantly faster when introduced to the epithelium-lined channel of the FT Chips and this increased significantly further when the FT Chips were differentiated in the presence of E2 that induces the highest level of cilia formation (**Figure 2E**). Interestingly, single particle tracking of the 5µm beads revealed the opposite trend (**Figure 3C, Videos S2-4**), where the particles moved at a higher velocity on empty chips compared to the FT Chips lined by the differentiated ampulla epithelium (**Videos S5-6**). Bead movement patterns on the E2 treated FT Chips were also highly heterogeneous. Consistent with this observation, microscopic analysis of bead localization revealed that many beads were trapped within the tissue folds (**Figure 3D**) while other beads appear to adhere to cilia for multiple days (**Video S6**), being swayed back and forth, occasionally being catapulted over longer distances; this likely contributed to the relatively large spread of values (**Figure 3E**). It is also important to note that the velocity of both beads was reduced in the NH FT Chip, which is consistent with our finding that expression of mucins (e.g. MUC5B, OVGP1) was higher in these chips, which should significantly increase local viscosity (η).

To distinguish between passive Brownian diffusion and active, directed transport mechanisms within the FT Chip, we analyzed the time-averaged Mean Squared Displacement (MSD) of particle tracks. The MSD provides a statistical measure of the area explored by a particle over a time interval 𝜏 and typically follows a power-law relationship: *MSD*(𝜏) = 4𝐷𝜏^𝛼^(**Equation 2**). Here, the anomalous diffusion exponent 𝛼 serves as an indicator of the mode of transport (**Figure 3F**). An exponent of 𝛼 ≈ 1 indicates normal Brownian diffusion. An exponent of 𝛼 < 1 (subdiffusion) typifies motion hindered by spatial confinement, crowding, or trapping. Conversely, 𝛼 > 1 (superdiffusion) indicates directed motion or active transport, such as that driven by ciliary beating or fluid flow.

We examined the transport dynamics of two distinct particle sizes—5 µm (sperm-sized) and 100 µm (oocyte-sized) from **Figures 3B** and **3F** using this metric (**Figures 3G, H**) and observed a striking divergence in their behavior on the FT Chips based on their size alone. In empty chips, the 100 µm showed minimal displacement due to their large size and surface friction (𝛼 = 0.72, **Figure 3G**, dark green line). Yet, when placed on the epithelium, these large beads were accelerated. In both biological conditions, they exhibited superdiffusive, parabolic trajectories (𝛼 > 1), moving far faster than they did in the empty channels (**Figure C,** lighter green lines). In contrast, for the 5 µm beads, transport was significantly hindered by the presence of the FT epithelium. In empty chips, these beads exhibited near-ideal Brownian motion (𝛼 = 0.94, **Figure 3H**, dark red line). However, in the presence of tissue (No Hormone condition), the beads became effectively trapped, resulting in a flattened MSD curve and a drastic reduction in displacement (**Figure 3H, bright red line**). Even under the active “fertile window” conditions (2 nM E2), the motion remained suppressed relative to the empty control (**Figure 3H**, light red line).

To evaluate how the same epithelial architecture can act as a trap for small particles while simultaneously acting as an accelerator for large particles, we developed a mechanism-based computational model.

We hypothesized that this size-dependent selection is a physical consequence of the ciliary geometry. We simulated the cilia field as a dense forest of obstacles (**Figure 3I**) and modeled the interactions based purely on particle size relative to ciliary spacing.

For the large particles, the simulation confirms an “Active Surface” regime. Because the 100 µm beads are too large to fit between the cilia, they rest on the ciliary tips. In this state, the cilia do not act as obstacles but as active actuators. The simulation successfully reproduces the transition from slow, passive diffusion in empty channels to rapid, superdiffusive transport on the ciliary carpet (**Figure 3L**). Furthermore, the transport efficiency for large beads increases with ciliary density (**Figure 3J**), as a denser field provides more frequent propulsive “kicks” to the cargo.

Conversely, our simulations reveal that particles smaller than the inter-ciliary spacing penetrate the ciliary layer and become trapped. The model replicates the experimental data, showing that while “Empty” simulations are linear (**Figure 3M**, dark red line), the presence of ciliary obstacles suppresses the effective diffusion, matching the hindered behavior seen in experiments (**Figure 3M**, light red line). A parameter sweep confirms this mechanism: as ciliary density increases (spacing decreases), the effective diffusion coefficient drops precipitously due to steric hindrance (**Figure 3K**). Interestingly, the model predicts an “Exclusion Regime” at extremely high densities (spacing < 5 µm), where small particles might be forced to the surface, potentially restoring mobility.

Together, these data illustrate a size-dependent selection mechanism intrinsic to the fallopian tube architecture. The geometry of the epithelium naturally segregates transport modes: the inter-ciliary space acts as a diffusive trap for small, passive particles (mimicking the head of immotile sperm), while the ciliary surface acts as an active conveyor for large cargoes (mimicking the oocyte).

#### The Fallopian Tube Chip recreates a living sperm reservoir in vitro

In contrast to the Cumulus-Oocyte-Complex (C-O-C) and early embryos, spermatozoa are highly motile due to the presence of the flagellum, which enables them to translocate through the lower female reproductive tract, reach the FT, and fertilize the oocyte. However, how human spermatozoa interact with the human FT remains largely unknown due to the inaccessibility of this tissue for direct observation. To study this, we injected spermatozoa into the proximal part of the FT chips and assessed how sperm translocate towards the distal part of the Chip over time, mimicking the translocation of spermatozoa through the native FT following an intercourse (**Figure 4A**).

**Figure 4.**
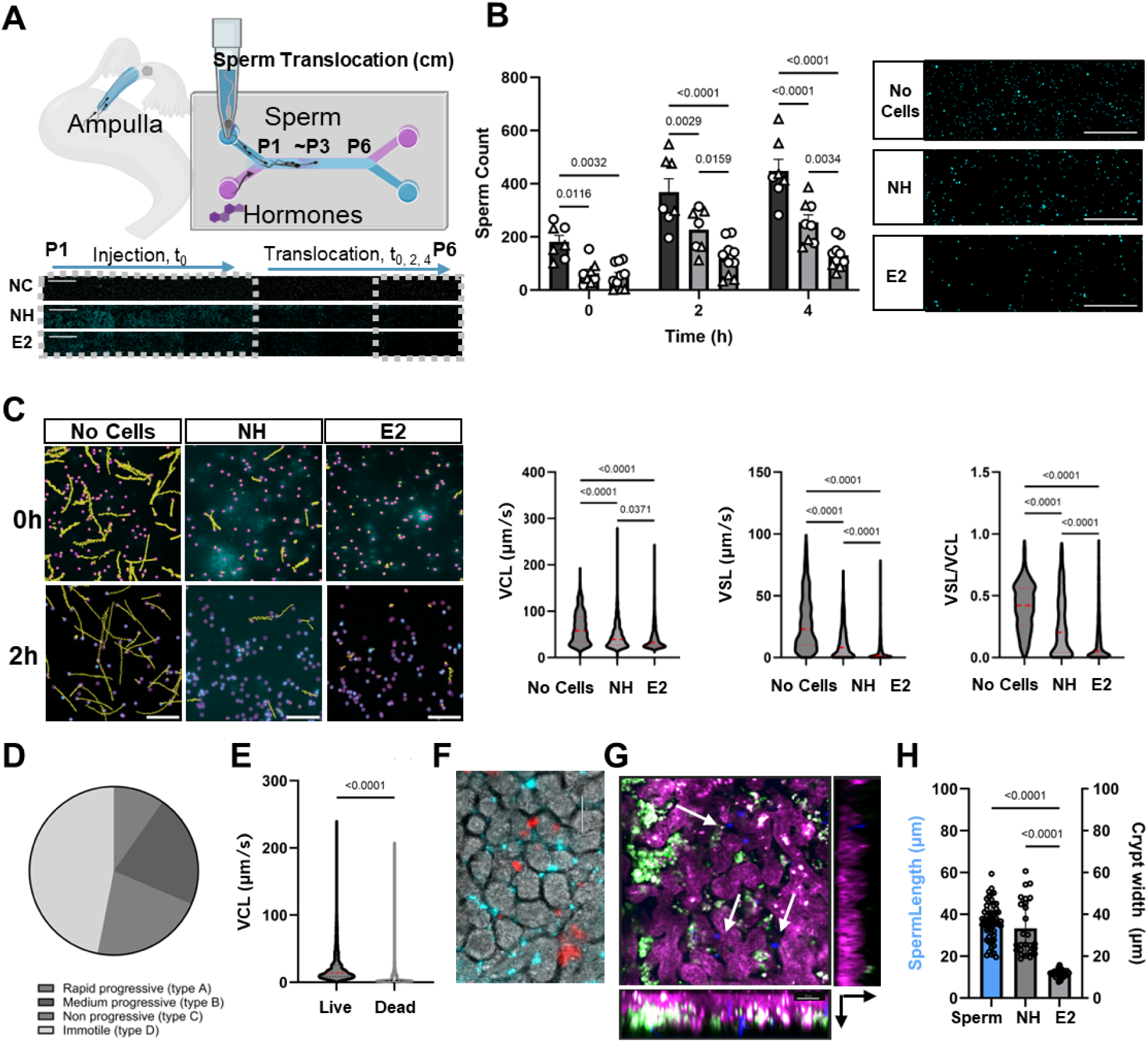
Follicular stage human Fallopian Tube Chips act like sperm reservoirs. (A) Diagram summarizing the sperm-FT Chip Translocation Assay workflow. Spermatozoa (nuclei, cyan) were introduced into the proximal region of the apical channel of the chip near the inlet ( blue arrow) and allowed to translocate through to P6 regions along the length of the 2 cm long channel (light blue arrow). Rectangular inset shows 6 images joined together visualizing the progressive reduction in the number of sperm appearing along the length of FT Chip. NC= no cells, NH = no hormones, E2 = 2nM Estradiol. Created in BioRender (B) Bar Graph depicting the number of sperm that transmigrated to the P6 region at the distal end of the channel immediately after injection (0 hr) versus after 2 or 4 hr of culture (N=7 chips from 2 epithelial donors and 6 human sperm donors pooled as 3 and 3 donors; data shown as Mean±SD; p values were determined using two-way ANOVA). Immunofluorescence images show sperm nuclei at P6 at 4 hours. Scale bar, 1000µm. (C) Representative images of sperm movement tracks (yellow) upon introduction on-chip (0 h) and 2h later (left) (Scale bar, 100µm). Violin plots at right depict corresponding curvilinear velocity (VCL), straight line velocity (VSL) and linearity of sperm movements measured using CASA at 2h (N=7, 8, and 10 chips from 2 epithelial donors and 6 human sperm donors; 535-1413 sperm were analyzed per group; p values were determined using one-way ANOVA). Corresponding Videos S.7-S.9 (D) Pie chart shows CASA-determined categories of FT E2 Chip released sperm at 72 hours after inoculation (red, rapid progressive; green, medium progressive; blue, non-progressive; grey, immotile; N=3 chips, and N=3 pooled sperm donors). Corresponding video S10(E) Violin plot depicting the curvilinear velocity (VCL) of matched live and dead spermatozoa on FT E2 Chips 2 hours after inoculation (N=3 chips, N=3 pooled sperm donors, N=1 epithelial donor, t-test). (F) A representative image of spermatozoa localizing in the FT E2 Chip folds. Sperm nuclei are shown in cyan and red dots correspond to 5µm beads. (Scale bar, 70µm). (G) Confocal reconstruction of cell tracker-stained FT E2 epithelium (magenta) and cilia (green) co-cultured with spermatozoa (blue). Live imaging. (Scale bar 50µm) (H) A Bar graph depicting the length of individual spermatozoa and the width of crypts on NH (no hormones) and E2 perfused FT Chips (N=2 biological epithelia donors, Mean±S.D., one-way ANOVA)

When we introduced donated sperm from 3 different pooled donors into the upper channel of an empty Organ Chip, we found that they were highly motile and could rapidly translocate to the distal end of the 2 cm long chip (**Figure 4B**). In contrast, the number of translocated spermatozoa was significantly lower when introduced into the epithelium-lined FT Chips cultured without steroid hormone (NH) when analyzed at either 15 minutes, 2 hours, or 4 hours. The number of translocated sperm was even significantly lower when introduced into the E2-stimulated “Fertile Window” follicular stage FT Chips compared to the NH Chips (**Figure 4B**). Thus, the follicular phase-associated morphological and biochemical changes we described in **Figures 1-3** may support the concentration of sperm within a localized region, hence mimicking the formation of a sperm reservoir that has been observed in many species *in vivo*^3^but has not yet been conclusively confirmed in humans.

We then utilized single particle tracking of the spermatozoa stained with Hoechst nuclear dye to gain insights into the local sperm motility (**Figure 4C, Videos S7-9**). Sperm tracks were analyzed to compute Curvilinear Velocity (VCL), Straight Line Velocity (VSL) and Linearity (VSL divided by VCL) of sperm movement, with Linearity being the most indicative of progressive translocation of the sperm through the FT. Spermatozoa in empty Organ Chips moved significantly faster (exhibited the highest VCL, VSL and Linearity) compared to when they were introduced to epithelium-lined FT Chips cultured without hormones. While the movement of the spermatozoa was slower in both the NH and E2 FT Chips, all parameters and especially the VSL and Linearity values were significantly lower in the follicular stage FT Chips that were exposed to E2. Intriguingly, this decrease in motility parameters was entirely due to the epithelial microenvironment and not caused by loss of sperm viability because when we physically cut open FT chips containing these stationary living sperm after 72 hours on the chip and released them into the culture medium dish, the freed sperm instantaneously restored their rapid movements (**Figure 4D, Supplementary Figure 7 and Video S10).** Despite the slower sperm movement in the E2-treated FT chips, the sperm also still remained motile locally as they displayed a VCL that was significantly higher compared to non-viable (95% CASA immotile) spermatozoa from matched donors (**Figure 4E**). Indeed, live microscopic visualization of Hoechst-stained spermatozoa together with the FT E2 Chip epithelia revealed that the sperm localize within the abundant tissue folds (**Figure 4F**). Interestingly, however, live confocal imaging did not suggest that the sperm bind directly to the cilia as seen in other species, and neither did we observe this type of interaction when f sperm were co-cultured with ciliated epithelial cells that were dissociated from FT E2 Chips (**Figure 4G**). Taken together, the decrease in sperm translocation and Linear movement in the FT Chip model appears to be largely due to the physical confinement within tissue folds.

#### Functional evaluation of non-hormonal contraceptives on-chip

Given the ability of the human FT Chip to recreate the ampulla microenvironment and support analysis of living human sperm movement, we leveraged this functionality to explore if we could use this as a preclinical model to test the safety and efficacy of non-hormonal contraceptives *in vitro*. We tested the FDA-approved contraceptive Nonoxynol 9 (N9) and a recently identified soluble adenylyl cyclase inhibitor, TDI-11861,^33^ that inhibits sperm capacitation and hyperactivation, which are required for fertilization^34^. As N9 is administered intravaginally in patients and TDI-11861 is designed to be given systemically, we introduced N9 (2%) through the apical epithelium-lined channel and TDI-11861 (3 μM) via the basal channel (**Figure 5A**) to evaluate their efficacy and safety. The membrane-disruptive agent N9 was used as a positive control to enable comparison to other contraceptive modalities^35^. Morphological analysis using DIC (**Figure 5B top**) and live imaging of beating cilia (**Figure 5B bottom)** with subsequent cilia area segmentation were used to evaluate the effect of these contraceptives on FT Chip structure and function. While the treatment with N9 resulted in a complete regression of tissue folds (**Figure 5B**), TDI-11861 exposure had no obvious effect on fold width or morphology (**Figure 5C**). Similarly, treatment with N9 resulted in a complete loss of ciliated cells while both the number of ciliated cells and cilia beating frequency remained unaffected after 24 hours of basal exposure to TDI-11861 (**Figure 5D**). Another central unanswered question in this field is whether systemically administered non-hormonal contraceptives can reach the luminal fluids of the FT or whether the FT epithelium is impenetrable to therapeutic compounds due to the anatomical inaccessibility of this compartment. This is especially difficult to ascertain in animal models due to the extremely small size of the FT in mice^36^. As a proof of concept, we perfused the FT Chips basally with the 3µM TDI-11861 for 24 hours while collecting the medium outflow from the apical channel. Automated Computer-Assisted Semen Analysis (CASA) software analysis confirmed that the outflows of TDI-11861-treated chips significantly inhibited the rapid progressive motility of fresh human spermatozoa, including reducing the percentage of rapid progressive spermatozoa as well as the VCL and VSL of motile spermatozoa, when the measurements were carried out using CASA standard slides (**Figure 5E**). These results are in line with the known effects of this compound, and we observed a similar level of inhibition as that displayed by N9 which disrupts the sperm lipid membranes.

**Figure 5.**
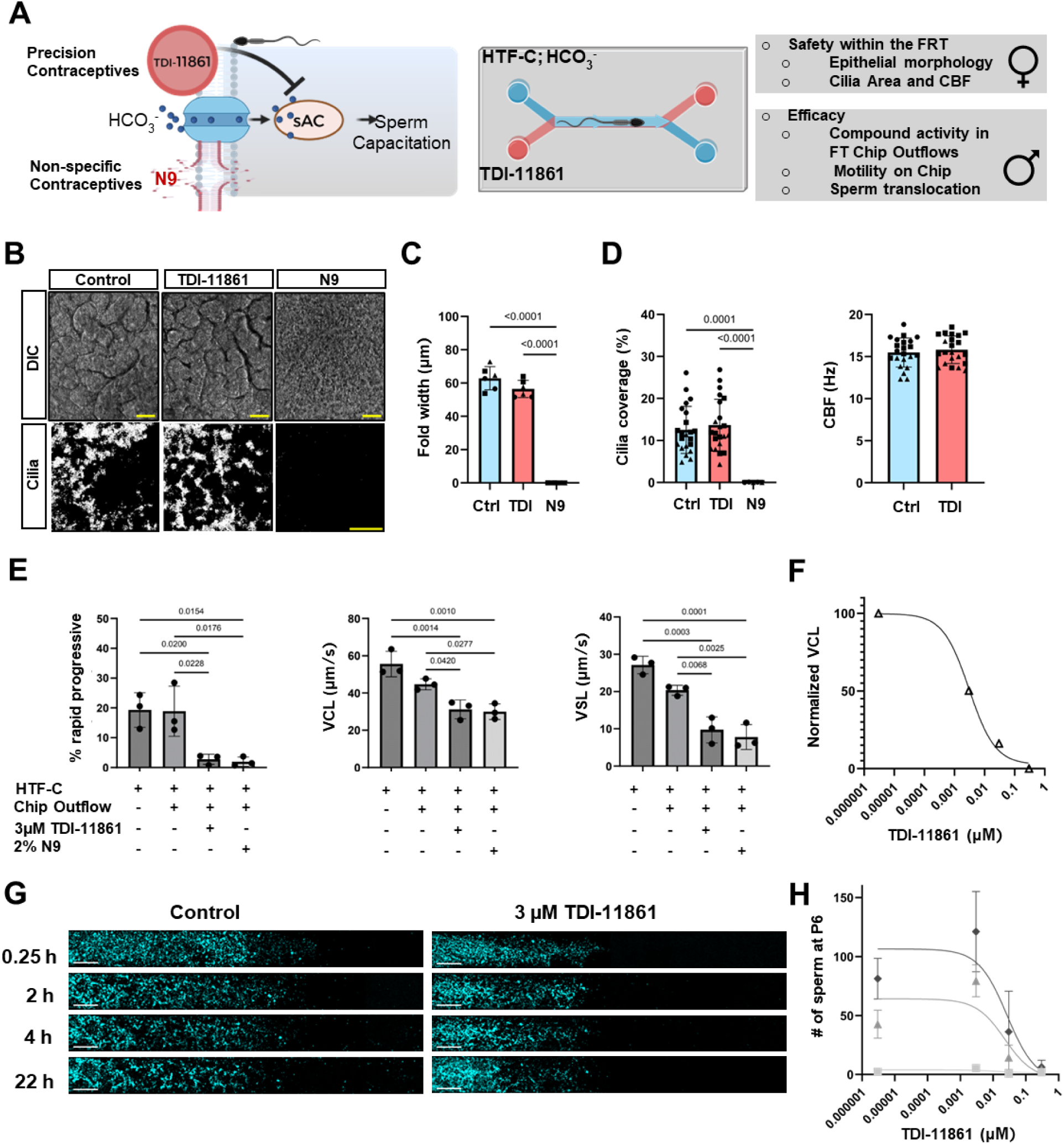
Fallopian Tube Chips enable functional evaluation of contraceptive efficacy and safety. (A) Diagram at left depicts the mode of action of the non-hormonal contraceptive TDI-11861 and the locally acting contraceptive Nonoxynol-9 (N9). The diagram on the right shows the contraceptive analysis workflow for analysis of the systemically administered TDI-11861, including the establishment of a relevant ionic environment in the apical channel and perfusion of the contraceptive through the basal channel. Created in BioRender. (B) DIC microscopic images (top) and visualization of beating cilia (bottom) on epithelium within E2-treated FT Chips that were either untreated for 24 hours (control) or exposed to either TDI-11861 (3µM) or N9 (2%) (Scale bar, 50µm). (C) Graph depicting changes in the width of the epithelial tissue folds 24 hours after continuous drug exposure. (N= 1 experiment, N= 3 chips, 2 measurements per chip; Mean ±SD, one-way ANOVA, Ctrl=untreated control; TDI=TDI-11861) (D) Graph depicting cilia coverage areas and cilia beat frequencies 24 hours after drug exposure (Data from 2 experiments, N= 22, 22, and 4 chips, Mean ±SD, one-way ANOVA, no significant difference was detected using a t-test). (E) Movement parameters determined by CASA for sperm treated with outflows from untreated chips (Con) or chips treated with N9 or TDI-11861 (data shown as Mean±SD are from 1 experiment; N=3 chips per group; N=3 pooled sperm donors; significance was determined using one-way ANOVA, VCL=curvilinear velocity, VSL= straight line velocity). (F) Graph depicting mean values and nonlinear fit shows dose-dependent effects of TDI-11861 on the normalized VCL for sperm 2 h after introduction into the chip. (Each dot represents the mean value of 400-1200 sperm tracks acquired across n=3-4 chips with sperm pooled from 3 human donors). (G) Tiled fluorescence microscopic images taken across the entire length of the chip depicting sperm (cyan) translocation after 0.25, 2, 4 and 22h (H). Graph shows dose-dependent inhibition of sperm translocation to the farthest P6 region of the channel at 0 (▪), 2 (▴) and 4(♦) hours after their introduction in chips perfused basally with 0, 0.003, 0.03 and 0.3µM TDI-11861 (Each symbol represents an average value for 3-4 FT Chips; mean±SD; nonlinear fit).

Importantly, the FT Chip also enabled direct monitoring of drug efficacy on-chip. Continuous basal perfusion of TDI-11861 at 0, 0.003, 0.03 and 0.3 μM inhibited sperm VCL in a dose-dependent manner when analyzed 2 hours after sperm introduction (**Figure 5F**). Images of sperm migrating over time in FT E2 Chips also confirmed that local sperm movement was inhibited by the sAC inhibitor TDI-11861 (**Figure 5G**). Finally, using the newly established functionally and physiologically relevant sperm translocation assay we developed (**Figure 4A**), we found that the TDI-11861 consistently acted in a dose-dependent manner to prevent translocation of spermatozoa along the length of the ∼2 cm long FT Chip (**Figure 5H**).

## DISCUSSION

In this study, we developed a microfluidic human FT Chip that recapitulates the structural, hormonal, and functional dynamics of the ampulla region of the human fallopian tube. This platform supports long-term epithelial differentiation under continuous flow, co-culture with human sperm, and steroid hormone perfusion, establishing a physiologically relevant model for both studying reproductive biology and carrying out preclinical testing in a human organ-relevant context.

Unlike conventional organoid systems^10,37^, the FT Chip reproducibly generates a folded epithelial architecture which is characteristic of the native fallopian tube. Importantly, matured FT Chips retained a balance of proliferative, secretory and ciliated phenotypes even at day 16, reflecting native tissue composition^11,38^.

In response to exposure to a high dose of estradiol to mimic the follicular phase of the menstrual cycle, the FT Chip exhibited marked increases in epithelial fold height, ciliation, and marked change in protein expression, including expression of a key estradiol regulated protein OVGP1^16^ and previously unappreciated proteins related to spermatozoa function. To our knowledge, this is the first demonstration of this level of menstrual cycle-specific hormone responsiveness in an *in vitro* FT model. The pronounced hormone responsiveness observed on-chip compared to in organoid culture^39^ likely results from sustained microfluidic flow exposure, which better supports lineage maturation than static cultures and leads to a distinct local availability of Wnt agonists and antagonists^10,40^. Similar effects of fluid flow on epithelial histodifferentiation have been observed in human Intestine Chips where dynamic medium flow in the basal channel is required for formation of folded intestinal villi.^41^.

Interestingly, the FT Chip revealed a dual functional role of estradiol in reproductive physiology as E2-driven changes in epithelial differentiation and ciliogenesis promoted the transport of oocyte-sized beads, validating a phenomenon previously observed only in rodents^42^, while simultaneously restricting sperm translocation over centimeter-scale distances. This finding suggests that periovulatory estradiol both enhances oocyte transport and facilitates sperm entrapment within the ampulla, optimizing fertilization efficiency. However, while the human FT Chip recreates this type of sperm reservoir, individual sperm cells retain their ability to move locally within the crypts between the abundant folds and retain their viability over multiple days. Similar to other primary human FT cell culture models^37^, the microfluidic FT Chip also prolonged sperm viability for at least 72 hours, underscoring its physiological fidelity. We speculate that in addition to biochemical changes induced by E2, the increase in the abundance and height of the epithelial folds and cilia may play a central role in creating a sperm reservoir by creating a topography that can trap sperm locally just before ovulation. These findings contextualize and align with results obtained using acellular patterned microfluidic channels^30,43^.

In 2019, around 162.9 million women lacked access to the contraception they needed^44^. While a broader option of non-hormonal contraceptive products could alleviate this needs gap, evaluating the safety and efficacy of novel precision contraceptive compounds has been a significant challenge due to the lack of suitable pre-clinical models^45^. The human FT Chip consists of two separately perfusable apical and basal channels, making it possible to systematically expose the tissues to both hormones and drugs/compounds using physiologically relevant routes of administration (i.e., intravaginal or systemic, respectively). The FT Chip also enables pharmacological testing within the physical context of a living human reproductive tract tissue. Using this model, we demonstrated that a recently proposed non-hormonal contraceptive that was shown to be effective in male mice^33^—the soluble adenylyl cyclase inhibitor TDI-11861—is also highly effective at inhibiting sperm motility in a human model of the FT without causing injury to the tissue. These human-relevant data are important because TDI-11861 is promising as a female-controlled contraceptive in line with global health female and maternal health priorities^46^.

Still, the current human FT Chip has some limitations. Several potentially critical stimuli were not incorporated into this model. For example, cyclic contractions of the fallopian tube have been suggested to contribute to sperm translocation within the female reproductive tract. While we did not explore this in the present study, our chips are able to exert cyclic mechanical forces that rhythmically deform the engineered tissue-tissue interface on-chip, and thus, this can be explored in future studies. Future studies may also benefit from extending this approach other anatomical parts of the fallopian tube beyond the ampulla, including the Isthmus. Another challenge for interpretation of the significance of this model is a lack of data relating to characterization of the human fallopian tube *in vivo*. However, it is precisely because it is so difficult characterize the human fallopian tube in vivo that the human FT Chip can be so valuable for studies on human fertilization and reproductive biology.

By integrating male and female cell reproductive interactions in a single human model, this Organ Chip platform addresses a major bottleneck in contraceptive development. Its scalability and alignment with the FDA Modernization Act^47^ further position the human FT Chip as a powerful alternative to animal models for advancing reproductive health research. This preclinical model may be used to complement the clinically cumbersome workflow of evaluating the safety and efficacy, which currently contributes to the significant bottleneck of developing non-hormonal contraceptives to address maternal health in developing countries. In the future, this system could be used to better understand infertility and inform the development of alternatives to IVF, a procedure currently inaccessible to half of the world. Taken together, the FT Chip provides an experimental platform that can be used to gain new insights into the human fallopian tube’s role in reproductive physiology, contributors to both male and female infertility, and serve as a preclinical model for evaluating the efficacy and safety of contraceptive strategies that can impact millions worldwide.

## RESOURCE AVAILABILITY

### Lead contact

Further information and requests for resources and reagents should be directed to and will be fulfilled by the lead contact, Donald Ingber.

### Materials availability

All non-cell materials can be purchased from commercial vendors.

### Data and code availability

- Bulk Proteomic Data have been deposited at MassIVE at MassIVE: MSV000099749and are publicly available as of the date of publication.
- Microscopy data reported in this paper will be shared by the lead contact upon request.
- Any additional information required to reanalyze the data reported in this paper is available from the lead contact upon request.

## ACKNOWLEDGMENTS

This work was supported by the Gates Foundation [INV-056272] and the Wyss Institute for Biologically Inspired Engineering at Harvard University. The authors thank Dr. Linu John for her input throughout the project execution and Dr. Janna C. Nawroth for her initial input on cilia analysis. The authors also thank Ms. Erin F Londregan for her help during the early stages of the project.

## AUTHOR CONTRIBUTIONS

Conceptualization, A.S. and D.E.I.; methodology, A.S, K.C., M.C; J.F, O.G., R.P., Investigation, A.S., K.C, M.C, J.F., S.T., N.B., A.G., M.M, M.L,; data analysis, A.S., M.C. J.F., computational modeling, D.G.; writing—original draft, A.S. with the inputs of D.G. on computational modeling; writing—review & editing all authors, A.S. and D.E.I.; funding acquisition, D.E.I.; resources, D.CH, J.P.; project management, A.J.; supervision. B.B. and D.E.I.

## DECLARATION OF GENERATIVE AI AND AI-ASSISTED TECHNOLOGIES

During the preparation of this work, the authors used ChatGPT (OpenAI) in order to conduct minor language editing. After using this tool/service, the authors reviewed and edited the content as needed and take full responsibility for the content of the published article.

## SUPPLEMENTAL INFORMATION

### LIST OF SUPPLEMENTARY VIDEOS

**Video S1.** Cilia on FT Chip and Organoid

**Video S2.** 5µm Bead Acellular

**Video S3.** 5µm Beads FT Chip No Hormones

**Video S4.** 5µm Beads FT Chip E2

**Video S5.** 5µm Beads with FT Chip Epithelia No Hormones

**Video S6.** 5µm Beads Trapped on Beating Cilia on FT Chip E2

**Video S7.** Sperm Hoechst on Acellular Chip

**Video S8.** Sperm Hoechst on FT Chip Estradiol

**Video S9.** Sperm Hoechst on FT Chip No Hormones

**Video S10.** Spermatozoa Released From the FT Chip E2 After 72 hours

## METHODS

### KEY RESOURCES TABLE

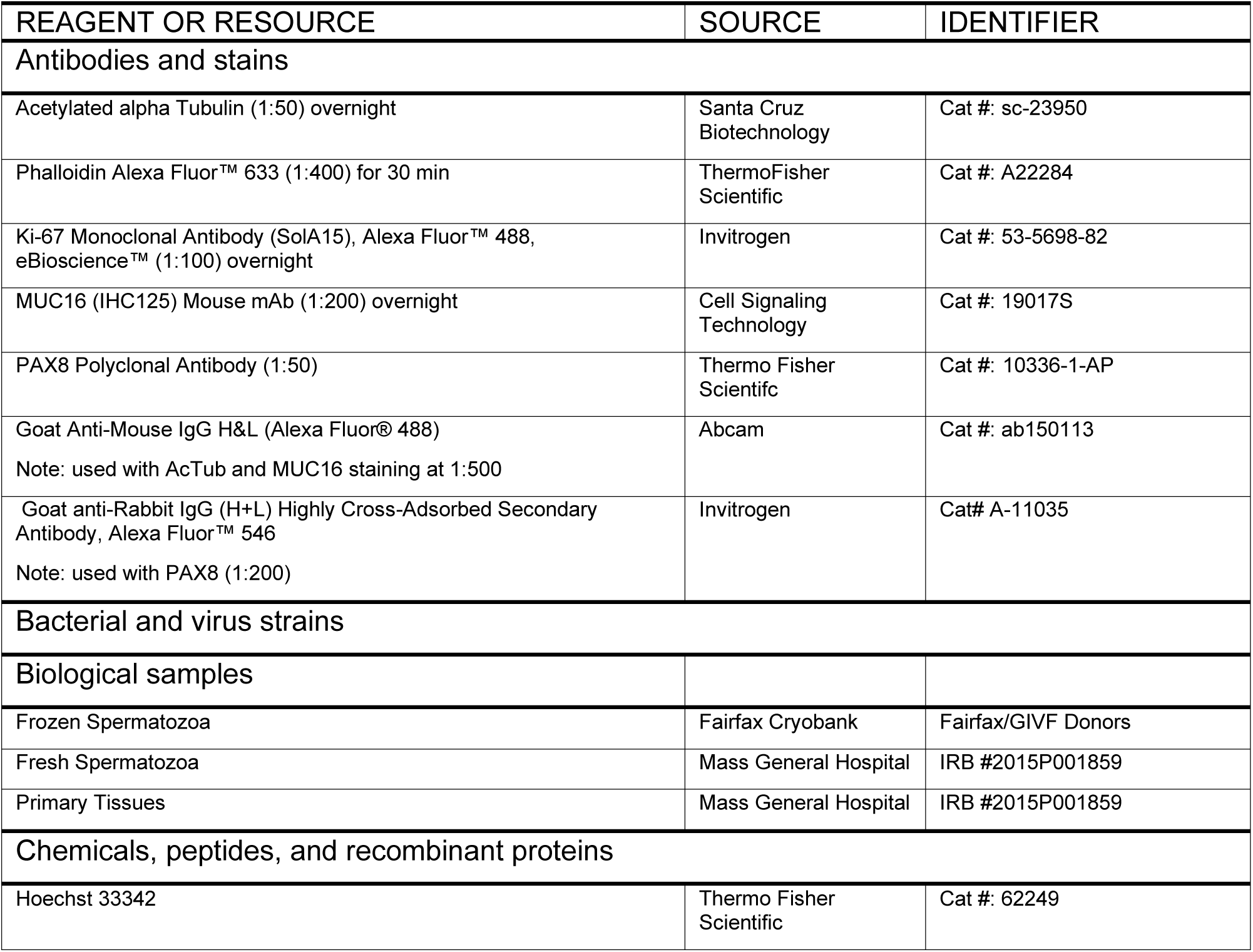
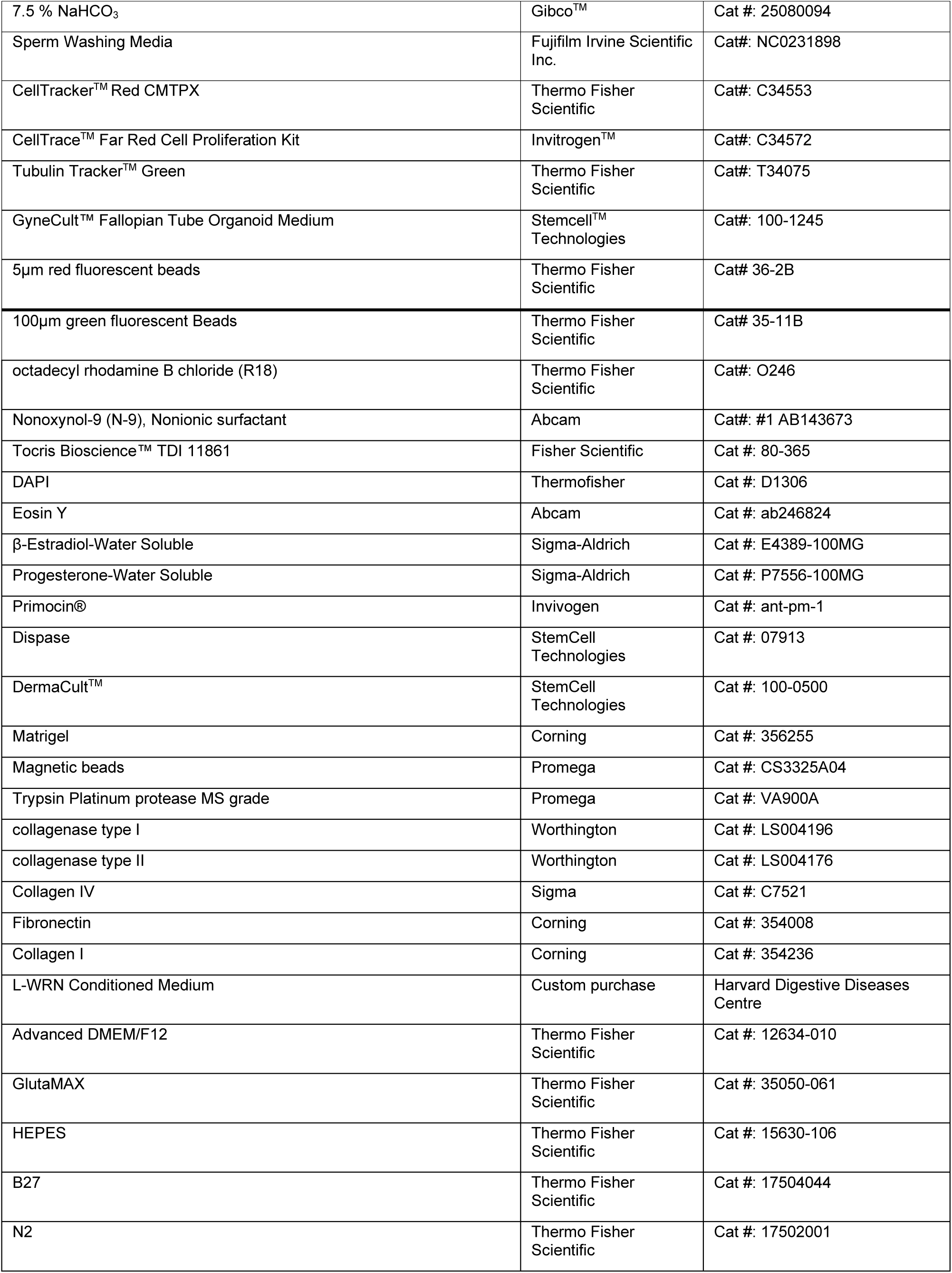
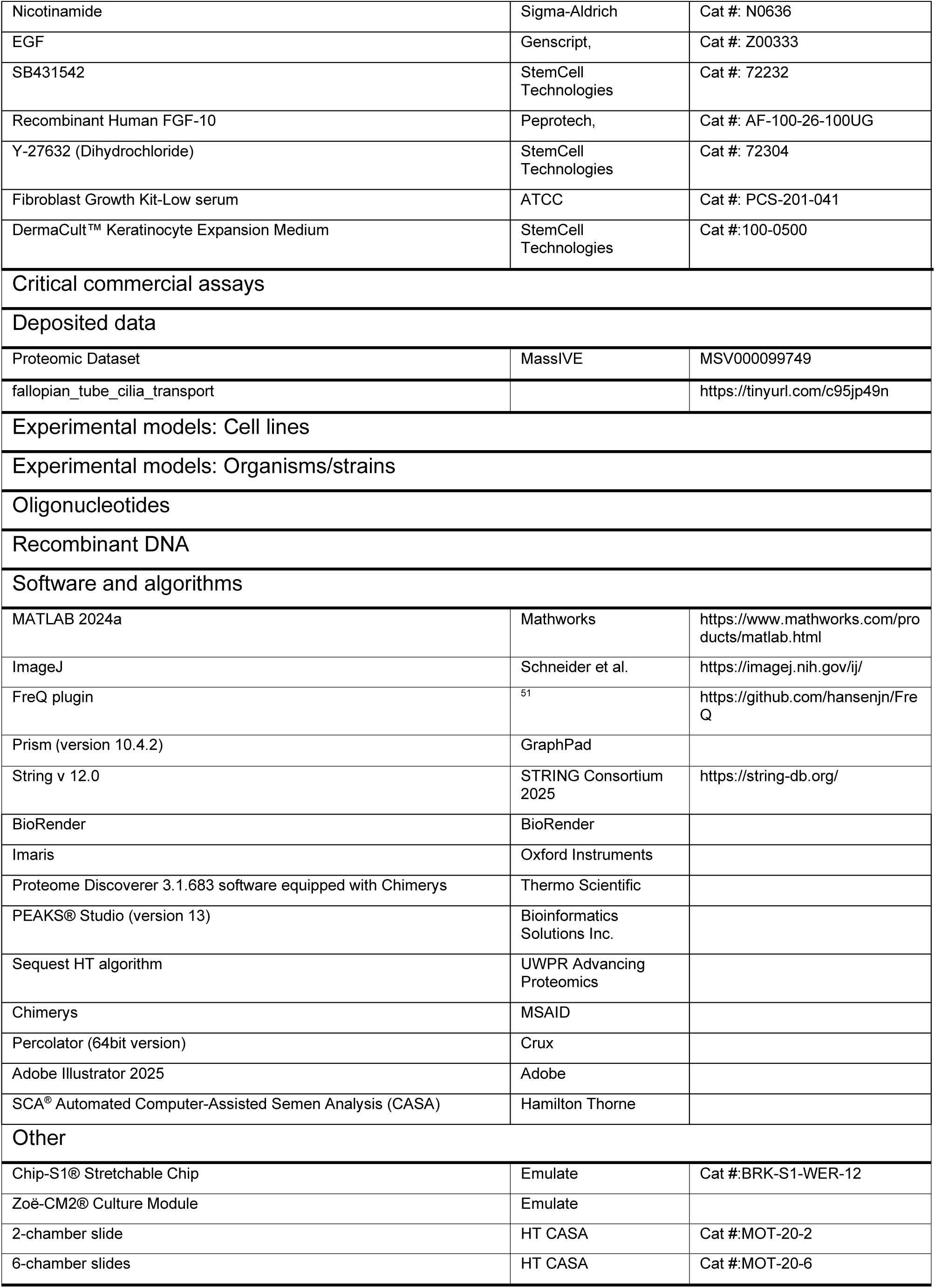

### EXPERIMENTAL MODEL AND STUDY PARTICIPANT DETAILS

The Ampulla Regions of the Fallopian Tube were anonymously collected from patients undergoing hysterectomy for benign conditions at the Massachusetts General Hospital under IRB-approved protocol #2015P001859. List of donors, including their age and reason for surgery, is provided in **Supplementary Table S1**. Fresh spermatozoa were acquired anonymously from donors undergoing semen analysis at the Massachusetts General Hospital under IRB-approved protocol #2015P001859. Only fertile donors, as determined by CASA analysis and adhering to the World Health Organization laboratory manual for semen analysis (6^th^ edition) were included in this study. Frozen sperm were acquired from the Fairfax Cryobank using sperm from fertile donors explicitly donated for research.

### METHOD DETAILS

#### Fallopian Tube Epithelial and Stromal Cell Isolation

The ampulla region of the Fallopian Tube was anonymously collected from patients undergoing hysterectomy for benign conditions (**Supplementary Table S1**) at the Massachusetts General Hospital under IRB-approved protocol #2015P001859. Explanted tissues were first washed in Phosphate Buffered Saline (PBS-/-; Gibco, 14190144) with Primocin (InvivoGen, ant-pm-1). Following the wash steps, the ampulla tissue sections were cut open longitudinally to expose mucosal folds/epithelial lumen. The processed tissues were then incubated in a solution containing 50% 5U/ml Dispase (STEMcell technologies, 07913) and 50% advanced DMEM/F12 (ThermoFisher Scientific, 12634010) with Primocin at 37 °C for 3 hours or at 4 °C overnight to enzymatically separate the epithelium from the stroma. The epithelium was then mechanically scraped away from the stroma using a Cell scraper (FALCON, 353085) and rinsed with PBS -/- between scrapings. The released epithelial cells were collected in a falcon tube, spun down at 300 g for 5 minutes, and the supernatant was discarded. Epithelial cells were subsequently treated with TrypLE™ Express Enzyme (1X) (ThermoFisher Scientific, 12604013) for 15 minutes at 37°C to dissociate cell clusters into single cells. Finally, the isolated fallopian tube epithelial cells were spun down at 300 g for 5 minutes, the supernatant was discarded, and cells were plated on T-75 or T-150 flasks depending on cell yield. Fallopian tube epithelial cells were expanded in an expansion medium containing 50% conditioned WRN medium, Advanced DMEM/F12 and other factors listed in (**Supplementary Table S3**) based on a previously published protocol and medium formulation^10^. After reaching 70-80% confluence cells were harvested and cryopreserved in FBS + 10% DMSO at passage P0. The stroma was digested using Collagenases Type I and II (Worthington, LS004196 & LS004176) at 37°C and the donor-matched stromal cells were expanded in DMEM (high glucose) supplemented with Primocin and 10% FBS (**Supplementary Table S2**)

#### Fallopian Tube Chip Culture

Two-channel microfluidic chips (Chip-S1®, Emulate) were surface-treated following the manufacturer’s protocol using ER-1 and ER-2 solutions (Emulate). The apical channel was then coated with 500 μg/ml collagen IV dissolved in 0.25% Acetic Acid (v/v) and the basal channel with 200 μg/mL rat tail collagen type I and 30 μg/mL fibronectin overnight at 37°C as previously described^48^. Both channels were washed twice with HBSS++ the next day. The apical channel was filled with fibroblast expansion medium, and the basal channel was then seeded with 50µl of primary ampulla fibroblasts (6 x 10^5^ cells/ml, passage P2). Chips were inverted for 2 hours to allow for fibroblast adhesion to the porous membrane rather than the bottom of the chip. One day later, the apical channel was seeded with 40µL of primary ampulla epithelial cells at passage P1 (1-2 x 10^6^ cells/ml) which were allowed to adhere overnight. Both cell types were seeded in their 2D expansion media. Finally, the chips were connected to flow the next day using Pods (Emulate) and Zoe Instrument (Emulate) and perfused at the rate 30µl/hr through both the apical and basal channels. The dynamic chip culture was divided into two stages: (i) expansion and (ii) differentiation. To achieve cell expansion, the basal channel was perfused with 50% defined low serum fibroblast medium (**Supplementary Table S4, S6**) and 50% WRN-medium (without Y27632) while the apical channel was perfused with 50% DermaCult^TM^ (StemCell Technologies, 100-0500) and supplemented with Hydrocrotisone, and WRN-medium (all components) (**Supplementary Table S5**) for three days. To promote differentiation, the growth factor-rich apical medium was replaced with a minimal HTF-medium (FUJIFILM Irvine Scientific, 9983) while the basal channel was perfused with GyneCult^TM^ (StemCell Technologies, 100-1244) supplemented with relevant custom hormones (**Supplementary Tables S7-8**). Standard differentiation was conducted in the presence of 2nM E2 unless otherwise stated (See section Hormonal Studies). During differentiation, the basal channel was continuously perfused at 45µl/hr and apical channel was perfused for 4 hours a day at 40µl/hr followed by 20 hours of stasis (0µl/hr).

#### Hormonal Studies

To evaluate the effect of hormones, the FT Chips were allowed to expand in the expansion media in the absence of exogenous hormones and subsequently perfused either with Gynecult^TM^ alone or supplemented with 2nM Estradiol E2 (Sigma-Aldrich, E4389-100MG) (E2 group) or 0.1nM Estradiol E2 and 1µM Progesterone P4 (Sigma-Aldrich, P7556-100MG) (E2 + P4 group) starting on day 5 of culture. The hormones were perfused through the basal channel at 45µL/hr throughout the differentiation phase to mimic systemic exposure.

#### Organoid culture

Fallopian tube epithelial cells were first expanded on T150 flasks in 2D expansion medium containing 50% conditioned WRN medium, Advanced DMEM/F12 and other factors following an established protocol^10^. After reaching 70-80% confluency, cells were harvested with trypLE and centrifuged at 300g for 5 minutes. Cell pellet was resuspended in Matrigel (Corning, 356255) at a density of 200,000 epithelial cells per millilitre. Subsequently, 20 uL Matrigel droplets (4000 cells per droplet) were seeded on 24-well plates. The well plate containing the Matrigel droplets was then incubated at 37°C for 20 minutes. Gynecult^TM^ medium (stemcell technologies) supplemented with 2nM E2 and Primocin® was then carefully added (as to not disrupt the Matrigel) to the well plate containing the Matrigel domes. Gynecult^TM^ media supplemented with Estradiol was refreshed every other day after initial seeding.

#### Relative epithelial height measurements

Relative epithelial layer height was measured manually using an inverted microscope (Axio Observer Z1; Zeiss) by reading and noting the z-coordinate of the in-focus plane of the PDMS membrane and the highest epithelial point at 10 random and independent locations covering the entire length of the chip. The readings were performed on at least 3 independent chips per condition. To eliminate bias, two researchers performed the readings.

#### Cilia Beating Frequency (CBF) Analysis: Microscopy Acquisition

Approximately 30 minutes before image acquisition, the inverted microscope (Axio Observer Z1; Zeiss) incubator was turned on and set to 37°C to allow sufficient time for thermal equilibration. Imaging was performed using a ZEISS Axio Observer Z1 with a Hamamatsu sCMOS camera and the ZEN Pro software. Two-second, 1024 x 1024 pixel videos were acquired at 40X/Ph2 in transmitted light (TL) Brightfield with a 5 ms exposure time (200 Hz) and 8–9V bulb voltage.

Three videos at independent locations were taken per chip.

#### Bead Movement Analysis

A bead master mix was prepared by combining 5 µm red fluorescent beads (Thermo Fisher Scientific, Cat# 36-2B) and 100 µm green fluorescent beads (Thermo Fisher Scientific, Cat# 35-11B) in capacitating sperm wash media (Sperm Washing Medium (FUJIFILM SKU:9983-100ML) supplemented with 25mM NaHCO3 (Gibco^TM^ Sodium Bicarbonate 7.5% solution, 25-080-094) per 10ml) and Primocin®. The red and green beads were diluted to final concentrations of 5,000,000 beads/mL and 10,000 beads/mL, respectively. The bead mixture was vortexed for 1 minute to ensure homogeneity and subsequently re-vortexed every 3–5 minutes to prevent settling of the larger green beads.

Immediately prior to injection, chips were disconnected from their pods. A total volume of 60 µL of the bead master mix was manually injected into the apical channel of each chip, filling the whole apical channel. During injection, pods were primed twice to ensure unobstructed flow.

After injection, approximately 5 µL droplets of the corresponding media were added to each chip’s inlet and outlet ports: Sperm Wash Media on the apical side and GyneCult™ media with relevant hormones on the basal side. Chips were then reconnected to their pods and equilibrated under basal flow (40 µL/hr) for 1 hour prior to imaging.

##### Microscopic Imaging

The microscope incubator was pre-heated to 37°C 30 minutes prior to imaging to ensure thermal equilibration. Imaging was performed using a ZEISS Axio Observer Z1 inverted microscope equipped with a Hamamatsu sCMOS camera and controlled via ZEN Pro software. Five-minute, 2048 x 2048 pixel videos were acquired at 40X under red fluorescence and 10X/1.6x Optovar under green fluorescence with a 100 ms exposure time and an acquisition rate of 20 frames per minute.

#### Sperm handling and preparation

##### Fresh Sperm Sample Preparation

Liquefied samples from three separate donors for each experiment repeat were obtained 1-5 hours after ejaculation on the day of each experiment from Massachusetts General Hospital under IRB-approved protocol #2015P001859. Sperm collected had total motility ≥45%, and the donors were categorized as fertile by healthcare professionals. Semen was separated from the seminal fluid and collected in separate 15-mL Falcon tubes by the donor and centrifuged at 300 x g for 5 minutes. The supernatant was discarded and 1 mL of Sperm Washing Medium (FUJIFILM SKU:9983-100ML) supplemented with 25mM NaHCO3 (Gibco^TM^ Sodium Bicarbonate 7.5% solution, 25-080-094) per 10ml, 1:500 Primocin® and 1:2000 Hoechst 33342 (Thermo Fisher, H1399) dye was gently layered over each sperm pellet. The 15-mL Falcon tubes were then incubated at 37 °C + 5% CO2 for 1 hour at a 45° angle to facilitate pellet swim-up of motile sperm. After swim-up the tubes were gently returned to an upright position and 75% of the supernatant from each donor was collected and combined into one tube for a total volume of ∼2.4 mL. An 8 µL sample from the combined supernatant tube was pipetted into a 20-um motility chamber, and an initial motility and concentration reading was taken using the SCA^®^ motility module with the SCA^®^ CASA system (Hamilton Thorne). The combined supernatant was then spun down at 300 x g for 5 minutes.

##### Frozen Sperm Sample Preparation and Assessment for Injection

Vials of cryopreserved human sperm from multiple healthy donors were obtained from Fairfax Cryobank and maintained on dry ice until use. For thawing, vials were placed in a 37°C water bath for 3–5 minutes. A 10 µL aliquot from each vial was applied to a counting slide (InCyto Cat#: DHC-N01) to assess motility under phase contrast microscopy. Samples with little or no motility were excluded from use. Per three viable sperm samples, a 15-mL Falcon tube was prepared with 4 mL of Sperm Washing Medium (FUJIFILM SKU:9983-100ML) supplemented with 5.5 µL of Hoechst dye and 11 µL of Primocin®. Once three samples were added the approximate volume should be 5.5 mL. The suspension(s) was incubated at room temperature for 10–15 minutes to allow for nuclear staining. Hoechst staining was monitored using fluorescence microscopy. After incubation, the suspension(s) were centrifuged at 300 × g for 5 minutes. Following centrifugation, the supernatant was aspirated, and 2 mL of Sperm Washing Medium was gently layered over the sperm pellet. The sample was incubated at 37 °C + 5% CO2 for 1 hour at a 45° angle to facilitate swim-up of motile sperm. After the 1-hour swim-up incubation, the tube was gently returned to an upright position. The upper supernatant layer, containing motile sperm, was carefully collected into a new tube, and its volume was recorded. A 10 µL sample from the supernatant was pipetted onto a Makler chamber which was then used to take an initial motility reading. The rest of the supernatant was then spun down at 300 x g for 5 minutes.

#### Sperm translocation and motility studies on the Chip

##### Sperm Collection and Resuspension for Sperm Translocation Assay

The supernatant (of either fresh or frozen sperm preps) was discarded and the pellet was then resuspended in the capacitating Sperm Washing Medium (Fisher Scientific, NC0231898, c-HTF) and 1:500 Primocin to a concentration of 20 million sperm/mL (**Supplementary Table S9**). The prepared sperm would then be diluted 1:1 to a concentration of 10 million/mL using collected and pooled apical outflows from chips separated by treatment conditions. Sperm were held in outflows for no longer than 30 minutes.

##### Sperm Injection into Organ Chips

All chips were disconnected from their pods prior to injection. A 10 µL volume of the appropriate sperm suspension, approximately 1/3.6 of the total apical channel volume determined to provide good signal to noise ratio in this assay, was injected into the apical outlet of each chip using a P100 pipette. While injecting, the pods were then primed twice. Following injection, droplets (∼ 5 uL) of corresponding media were applied to each chip’s inlet and outlet ports, and chips were reconnected to their pods. The basal & apical outlet reservoirs were aspirated, and imaging commenced immediately.

##### Sperm Collection, Resuspension, and Injection for Whole Chip Introduction

The supernatant (of either fresh or frozen sperm preps) was discarded and the pellet was then resuspended in capacitating Sperm Washing Medium (c-HTF) and 1:500 Primocin to a concentration of 5 million sperm/mL. All chips were disconnected from their pods prior to injection. The prepared sperm would then be diluted 1:1 to a concentration of 2.5 million/mL using collected and pooled apical outflows from chips separated by treatment condition. Sperm were held in outflows for no longer than 30 minutes. A 40 µL volume of sperm suspension was injected into the apical outlet of each chip. While injecting, the pods were then primed twice. Following injection, droplets (∼ 5 uL) of corresponding media (Sperm Wash Media apically and basal media basally) were applied to each chip’s inlet and outlet ports, and chips were reconnected to their pods. The basal & apical outlet reservoirs were aspirated, and imaging commenced immediately.

##### Sperm Translocation Imaging

Approximately 30 minutes prior to image acquisition, the microscope incubator was set to 37 °C to ensure thermal equilibration. Imaging was performed immediately following sperm introduction using a ZEISS Axio Observer Z1 equipped with a Hamamatsu sCMOS camera and controlled via ZEN Pro 2 software. Sperm were visualized using Hoechst fluorescence to target the blue-stained nuclei. Each chip was imaged using 8–12 tiles to capture the full length of the channel. Images were acquired at a resolution of 2048 × 2048 using a 5× objective with a 1.6× Optovar. One stitched tile per chip was retained for analysis.

#### Contraceptive Screening Studies

The apical medium in drug screening experiments consisted of capacitating HTF (c-HTF), prepared by supplementing HTF with sodium bicarbonate. The basal medium consisted of GyneCult^TM^ supplemented with E2. Both media contained Primocin. To evaluate the efficacy and safety of TDI-11861 (MecChem Express, HY-151798)(0; 0.003; 0.03, 0.3 and 3µM) and Nonoxynol-9(N-9, abcam, ab143673, 2%), TDI-11861 was perfused basally using continuous flow (45µl/hour) while 2% N-9 was introduced apically for 4-hours at 40µl/hour followed by 20 hours of stasis and this regime was repeated daily. The media were prepared and refreshed daily.

##### Effect of Contraceptives on FT Chip Epithelium Morphology and Ciliation

The chips were imaged 24-hours post first exposure to contraceptives (3µM TDI-11861, 2% N-9) using DIC imaging and video microscopy to determine the effect of the compounds on epithelial morphology and cilia beating frequency (CBF).

##### Evaluation of the Apical Outflow Sperm-inhibitory Activity

To determine whether TDI-11861(3µM) perfused basally can reach the apical lumen and can exert contraceptive effect in this compartment, the Apical outflows were collected 24-hours (4 hour apical flow, 20 hours of stasis, 40/40 µl/hour) after the contraceptives were introduced on the Chips. The inhibitory activity of the outflows (VCL, VSL) was evaluated using fresh sperm from 3 pooled donors in 2D using the CASA system. The sperm concentration in these studies was 1 million/mL and the samples were prepared by mixing 25µL sperm aliquots (c-HTF) with 75µl of apical outflows per well and assessed at 2 hours. Outflows from FT-Chips consisting of c-HTFC + Primocin® treated with 2% N-9 and from non-treated FT Chips were used as controls

##### Effect of TDI-11861 on spermatozoa on FT Chips

The sperm were injected on Chips 48-hours post first Fallopian Tube Chip drug exposure in a mixture of treatment group-matched fresh chip apical outflows and fresh c-HTF and sperm motility and translocation were measured at defined time intervals. Following sperm introduction, FT Chips were perfused continuously with the relevant TDI-11861 concentration through the basal channel at 45 µL/hr while the apical channel was kept static.

##### Media Preparation and Conditioning

For sperm pre-conditioning, media was flowed through each chip under normal apical/basal flow conditions (45/45 µL/hr) to accumulate sufficient apical outflow for sperm resuspension (∼100 µL per condition) for 1 hour.

#### Bulk Proteomics of the Fallopian Tube Epithelial Cells

##### Sample preparation for proteomics

###### Protein Extraction Protocol

Both FT Chips and 2D-adherent FT epithelial cells were washed with HBSS++ prior to digestion. Cell pellets, derived either from primary fallopian tube epithelial cells expanded in 2D or TrypLE Express (ThermoFisher Scientific)-cleaved epithelial cells from the apical channel of mature FT Chips, were frozen in PBS for proteomics. The thawed cells (30µL – 10ug of protein) were immediately submitted to lysis using 6M Guanidinium Chloride (spec) (1:1 v/v) and vortexing for 2 minutes.

###### Reduction/Alkylation

10μL of 10mM TCEP (tris(2-carboxyethyl)phosphine) in 50mM TEAB was added to each filter and samples were incubated for 45 minutes at 350 rpm and 25°C on Eppendorf ThermoMixer C. After reduction, 10μL of 10mM iodoacetamide in 50mM TEAB (Tetraethylammonium bromide) was added to the samples for alkylation, and they were incubated for 15 minutes at 350 rpm and 25°C.

###### SP3 Magnetic Bead Digestion

Protein samples (10 μg per sample) were processed using a modified SP3 (Single-Pot Solid-Phase-enhanced Sample Preparation) protocol to enable efficient protein binding, clean-up, and on-bead digestion.

#### Bead Preparation

Magnetic beads (Promega, cat. number CS3325A04) were vortexed thoroughly to ensure homogeneity, and 30 μL (500 μg) aliquots were transferred for each sample, in accordance with the vendor’s protocol. Beads were washed three times with high-purity water using a magnetic rack to separate beads and discard supernatant between washes.

#### Protein Binding

Reduced and alkylated protein samples were added to the washed beads. Ethanol was added to achieve an 80% final ethanol concentration. The mixture was vortexed for 30 seconds and incubated with agitation at 1,200 rpm for 20 minutes to promote protein binding to the beads. Beads were then captured on a magnetic rack, supernatants removed, and the bead-protein complexes were washed three times with 1 mL of 80% ethanol.

#### On-Bead Digestion

Proteins bound to beads were resuspended in 50 mM TEAB buffer and Trypsin Platinum protease MS grade (Promega, Ref VA900A) (100μg/mL) was added at a 1:50 enzyme-to-protein ratio to the samples. Digestion proceeded overnight at 40°C with agitation at 1,200 rpm. Post-digestion, beads were separated magnetically and the supernatant collected. Beads were then washed once with 50 mM TEAB for maximum collection of peptides, and the wash combined with the initial supernatant. The final peptide solution was dried and resuspended in 10 μL of Buffer A (0.1% formic acid in ultrapure HPLC grade water) for subsequent LC-MS/MS analysis.

#### Mass spectrometry analysis

Each sample was submitted for single LC-MS/MS experiment that was performed on a Q-Exactive Orbitrap (Thermo Scientific) equipped with EVOSEP ONE (EVOSEP) nanoHPLC pump. Peptides were separated onto a PepSep C18 15cmx150um, 1.9um analytical column (Bruker Daltonics GmbH & Co KG). Separation was achieved through a 44 min active reverse phase gradient with a 48 min total cycle time. Mobile phase A is 0.1% formic acid in water and mobile phase B is 0.1% formic acid in acetonitrile, and the flow rate is 500 nL/min. Electrospray ionization was enabled through applying a voltage of 2 kV using a PepSep electrode junction at the end of the analytical column and sprayed from stainless still PepSep emitter SS 30µm LJ (Bruker, MA). The Q-Exactive Orbitrap was acquired in data-dependent acquisition mode for the mass spectrometry methods. The mass spectrometry MS1 survey scan was performed across the scan range of 450 –900 m/z at a resolution of 30K, followed by the selection of the ten most intense ions (TOP10) ions, that were subjected to HCD MS2 event in the orbitrap part of the instrument. The fragment ion isolation width was set to 0.8 m/z, AGC was set to 3 x 10^6^, the maximum ion accumulation time in the C-trap was 150 ms, normalized collision energy was set to 34V and an activation time of 1 ms for each HCD MS2 scan. The resolution for MS2 scan is set to 70K.

##### Proteome data analysis

The raw data was submitted for analysis in Proteome Discoverer 3.1.683 (Thermo Scientific) software equipped with Chimerys. The assignment of MS/MS spectra was performed using the Sequest HT algorithm and Chimerys (MSAID, Germany) by searching the data against a protein sequence database including all entries from the Human Uniprot database (Homo Sapiens Human Uniprot) and other known contaminants such as human keratins and common lab contaminants. Sequest HT searches were performed using a 20 ppm precursor ion tolerance and requiring each peptide’s N-/C- termini to adhere with Trypsin protease specificity, while allowing up to two missed cleavages. 18-plex TMT tags on peptide N termini and lysine residues (+304.207146 Da) were set as static modifications and carbamidomethyl on cysteine amino acids (+57.021464 Da) while methionine oxidation (+15.99492 Da) was set as a variable modification. A MS2 spectra assignment false discovery rate (FDR) of 1% on protein level was achieved by applying the target-decoy database search. Filtering was performed using a Percolator (64bit version)^49^. For quantification, a 0.02 m/z window was centered on the theoretical m/z value of each the 18 reporter ions and the intensity of the signal was recorded closest to the theoretical m/z value. The reporter ion intensities were exported into a result file of Proteome Discoverer search engine as excel sheets. The total quantified signal intensities across all peptides were summed for each TMT channel, and all intensity values were adjusted to account for uneven TMT labeling and/or sample handling variance for each labeled channel.

#### MSD Simulations

Particle transport was modeled using an overdamped Langevin dynamics framework implemented in MATLAB 2024a. The simulation domain was populated with rigid, circular obstacles ($r = 0.8$ µm) distributed randomly to mimic the ciliary field, with an average inter-ciliary spacing of 10 µm. Particle trajectories were calculated with a time step of 0.05 s over a total duration of 60 seconds. The physical interaction regime was determined by the particle diameter relative to the ciliary spacing. For particles smaller than the gap threshold (5 µm), dynamics were governed by steric hindrance, where collisions with ciliary stems were resolved via displacement reflection and scaling. For particles larger than the gap threshold (100 µm), the regime shifted to surface transport; particles were modeled as floating above the obstacle layer, subject to a stochastic active bath consisting of a constant velocity magnitude and random orientation at each time step. Baseline passive diffusion coefficients were derived from the Stokes-Einstein equation assuming the viscosity of water at 298 K. The full code is deposited at https://tinyurl.com/c95jp49n.

#### Live-Cell and Cilia Staining

Live cilia were visualized using the Tubulin Tracker^TM^ Green (ThermoFisher Cat: T34075) at 1X concentration and the epithelial cells were visualized using the CellTrace^TM^ Far Red Cell Proliferation Kit (Invitrogen Cat: C34572) (10 µM). To stain the live FT Chips, the Chips were washed once with HBSS+/+ and subsequently incubated in a mixture of these two dyes for 30–45 minutes at 37°C and gently washed three times with HBSS+/+ to remove any unbound dye prior to being co-cultured with the spermatozoa.

#### Immunostaining

Mature fixed FT Chips (day 16) were stained with Acetylated alpha Tubulin (1:50) to visualize cilia, Phalloidin Alexa Fluor 633 (1:400) to visualize f-actin, Ki-67 Monoclonal Antibody (1:100) to visualize proliferating cells and MUC16 (IHC125) (1:200)and PAX8 (1:50) to visualize secretory lineage. Apical and basal channels of the chips were washed three times with PBS-/-(Gibco, 14190144) Both channels of the chip were fixed with 4% PFA (ThermoFisher, J61899.AP) for 1 hour at room temperature and subsequently washed three times with PBS. Prior to processing immunostaining, chips were stored at 4°C in PBS. Chips were either sectioned prior to staining (for sideview images) or were stained whole (top view images). Chip sectioning was performed as follows: excess PDMS was cut away from the channels using a blade, the chip was then cut in half transversely, mounted on the vibratome specimen holder (Leica VT 1000 S) and sectioned at a thickness of 200 µM. Fixed whole chips or sections were then permeabilized with 0.1 % Triton-X (SigmaAldrich, X100) in PBS for 20 minutes at room temperature, washed three times with PBS, then blocked with 1% BSA in PBS (blocking solution) for 45 minutes at room temperature. Chips were then incubated with primary antibodies (diluted in 1% BSA in PBS) at 4°C overnight. The next day, chips/sections were washed with PBS three times to remove unbound primary antibody. Subsequently, samples were incubated with secondary antibodies (diluted in 1% BSA in PBS) at room temperature for 1 hour. Chips/sections were then rinsed with PBS three times to remove unbound secondary antibody and stored in PBS at 4°C prior to imaging

#### Microscopic Imaging

Live phase contrast (organoids, chips) and fluorescence imaging (beads, spermatozoa) was conducted using Revolve (ECHO) microscope in the inverted mode. To visualize the Chip morphology, differential interference contrast (DIC) microscopic imaging was utilized on both fixed (4% paraformaldehyde) and live cells using an inverted microscope (ZEISS Axio Observer Z1). Immunostained samples (sections and whole chips) were imaged on a Stellaris confocal microscope (Leica). Live imaging of cilia beating frequency and sperm motility is described separately. Images were visualized using FIJI (Image J)^50^.

### QUANTIFICATION AND STATISTICAL ANALYSIS

#### Ciliary Beat Frequency (CBF) Video Analysis

Following acquisition, videos were exported for offline analysis. Ciliary beat frequency was quantified using the FreQ plugin^51^ for FIJI/ImageJ, which is publicly available with instructions for download and installation at https://github.com/hansenjn/FreQ.

For analysis, videos in .CZI format were imported directly into FIJI/ImageJ and converted to .tif files. No other pre-processing was performed. Once the videos were loaded into FIJI, the plugin was accessed by navigating to the “Plugins” menu, selecting “JNH,” and then launching “FreQ.” Within the FreQ interface, default settings were reviewed, and the recording frequency was adjusted from 2000 Hz to 200 Hz and the box for complex maps was checked. The analysis was then executed.

The primary frequency and primary power .txt files that FreQ generates were run through a MATLAB script to calculate average and median values for both. Additionally, the corrected signal area .tif files that FreQ output were imported back into imageJ/FIJI, thresholded, and percentage area for each video was calculated. The outputs were averaged over the 3 videos acquired per chip.

#### Epithelial Fold Width and Crypt Width Measurements

Epithelial fold and crypt width were measured manually in imageJ using 2-5 DIC images/chip. 10 measurements were taken per image and the average fold or crypt width of the 10 measurements of a single image was used. Each data point represents the average fold or crypt width per one image. (N= 8 Chips; n= 25-26 images; results from two experiments; p-values determined by one-way ANOVA).

#### Quantification of Bead Movement on the Chip

Bead tracking was conducted using the TrackMate plugin^52^ in FIJI/ImageJ^50^. Spot detection was performed using the LoG (Laplacian of Gaussian) detector with an estimated blob diameter of 7 µm and a quality threshold of 50 for the 5 µm red fluorescent beads. The same method of detection was used for the 100 µm green, fluorescent beads, but with an estimated blob diameter of 100 µm and a quality threshold of one. No filters were applied to detected spots.

Tracks were created using the LAP tracker with the following settings: Frame-to-frame linking max distance was set to 15 µm, Gap-closing max distance was set to 15 µm, Gap-closing max frame gap was set to 2 frames, Track segment splitting and track merging were both disabled. The same settings were used for both bead types. Tracks with a duration < 6 frames were excluded from analysis. The remaining track data were exported as CSV files and analyzed downstream

#### Sperm Analysis on the Chip

Following acquisition, images were imported into FIJI (ImageJ) for processing. Each stitched tile was straightened and cropped to exclude regions where the apical and basal channels diverged. A custom macro was then applied to each tile to automate background subtraction, segment into six equal zones, color balance, threshold, and quantify using the “Analyze Particles” function. Sperm counts were assessed by position along the chip to evaluate distribution and translocation efficiency in response to treatment with TDI-11861.

*Sperm Motility Imaging & Analysis*: Videos were captured immediately following the acquisition of the tiling images using the same microscope and fluorescent channel as detailed previously. Two-second videos were acquired at a speed of 50 FPS (20 ms exposure) at a resolution of 2048 x 2048 using a 20x objective with a 1.0x Optovar. Three videos per chip were taken and retained for analysis.

To analyze the acquired videos a new plug-in was coded that analyzed multiple videos simultaneously using the Track Mate plug-in. The plugin first subtracted any background present and enhanced contrast of all frames. The LoG spot detector was used with an estimated object diameter of 7 µm and a quality threshold of 0.357. The following settings were used for the LAP tracker:

A track duration of 0.3 seconds (6 frames) was applied to filter out any false readings. The created plugin ran these settings and created an excel sheet for each video that contained track info for each detected sperm. The excels were then used for further downstream analysis.

#### Sperm Analysis in the Outflows

Sperm motility was evaluated using a Hamilton Thorne Computer-Assisted Sperm Analysis (CASA) system (Sperm Class Analyzer-SCA, software version 6.7.30; Hamilton-Thorne Research, USA) equipped with a Nikon Eclipse Ci-L microscope (Nikon, Japan) and a Basler Ace ACA1300-200UC camera. The CASA system enabled objective quantification of sperm movement by capturing digital images and generating individual sperm trajectories for subsequent kinematic analysis. Motility parameters were assessed according to the World Health Organization Laboratory Manual for the Examination and Processing of Human Semen, 6th Edition (WHO6)^53^. The following parameters were recorded: total motility (%), sperm concentration (*10^6^/ml), progressive motility (%), average path velocity (VAP, μm/s), straight-line velocity (VSL, μm/s), curvilinear velocity (VCL, μm/s), straightness (STR, VSL/VAP, %) and linearity (LIN, VSL/VCL, %).

#### Analysis of Proteomic Differential Expression using WASP

##### Database Search and Bioinformatics Differential Expression

Peptide Spectrum Matches (PSMs) were aggregated to the peptide level via PEAKS (version 13). Next, peptide level abundances were normalized using the Trimmed Mean of M values (TMM) method, null values imputed by left-tail sampling, log2 transformed, and aggregated to the protein level by the mean of the top 3 most intense peptides. Differential abundance was conducted by limma (version 3.21) in R (version 4.5.0), and proteins with |log2 fold change > 0.58| and p-value < 0.05 were classified as differentially abundant.

The output Limma_reuslts-isoforms matrix was filtered using the following criteria: adjusted P value <0.1 (comparison of the effect of hormones) and adjusted P value <0.01 (comparison of the 2D cells and Chips). The upregulated and downregulated targets below these adjusted P values were separately analyzed using STRING v12.0 to identify enriched biological processes. The most prominent processes ranked according to the parameter ‘Signal’, a weighted harmonic mean between the observed/expected ration and -log(False Discovery Rate or FDR) were visualized using GraphPad Prism.

To identify the number of upregulated and downregulated proteins in E2 and E2+P4 group compared to controls, the number of proteins Limma_reuslts-isoforms matrix with p value below p<0.1 were quantified and plotted.

#### Statistical analysis

Data were analyzed and visualized using GraphPad Prism (version 10.4.2) or MATLAB 2024a unless otherwise stated. The statistical tests included Student’s t-test, one-way ANOVA and two-way ANOVA. The significance levels were determined as: *p<0.05, **p<0.01, and ***p<0.001 and where possible exact significance levels are included. Dose-response to TDI-11861 curve was generated using the nonlinear fit function [Inhibitor] vs. response (three parameters) in GraphPad Prism on a normalized (VCL) or original dataset. Chip Quality Control Analysis was also used to ensure reproducibility of results. Chips were regularly checked for change in media color and blunting of folds or flow disruption in the microfluidic devices and chips that exhibited mixed flow, detached epithelia or excessive bubbles were excluded. We note, however, that such exclusion events were rare for this model.

## Supplementary Tables

**Supplementary Table S1:** List of Donors.

| Donor | Donor Age and Sex | Reason for surgery | Comments |
| --- | --- | --- | --- |
| 1 | 37yo Female | Menorrhagia | Cilia observed during epithelial cell isolation |
| 2 | 35yo Female | Voluntary sterilization | Cilia observed during epithelial cell isolation |
| 3 | 33yo Female | Adenomyosis | Cilia not observed during epithelial cell isolation |

### Expansion on 2D culture flasks

**Supplementary Table S2:** Fibroblast Expansion Medium.

| Component | Content | Catalog # |
| --- | --- | --- |
| DMEM, high glucose, GlutaMAX | 90% | Thermo Fisher Scientific, 10-566-016 |
| Fetal Bovine Serum | 10% | Thermo Fisher Scientific, 10082-147 |

**Supplementary Table S3:**
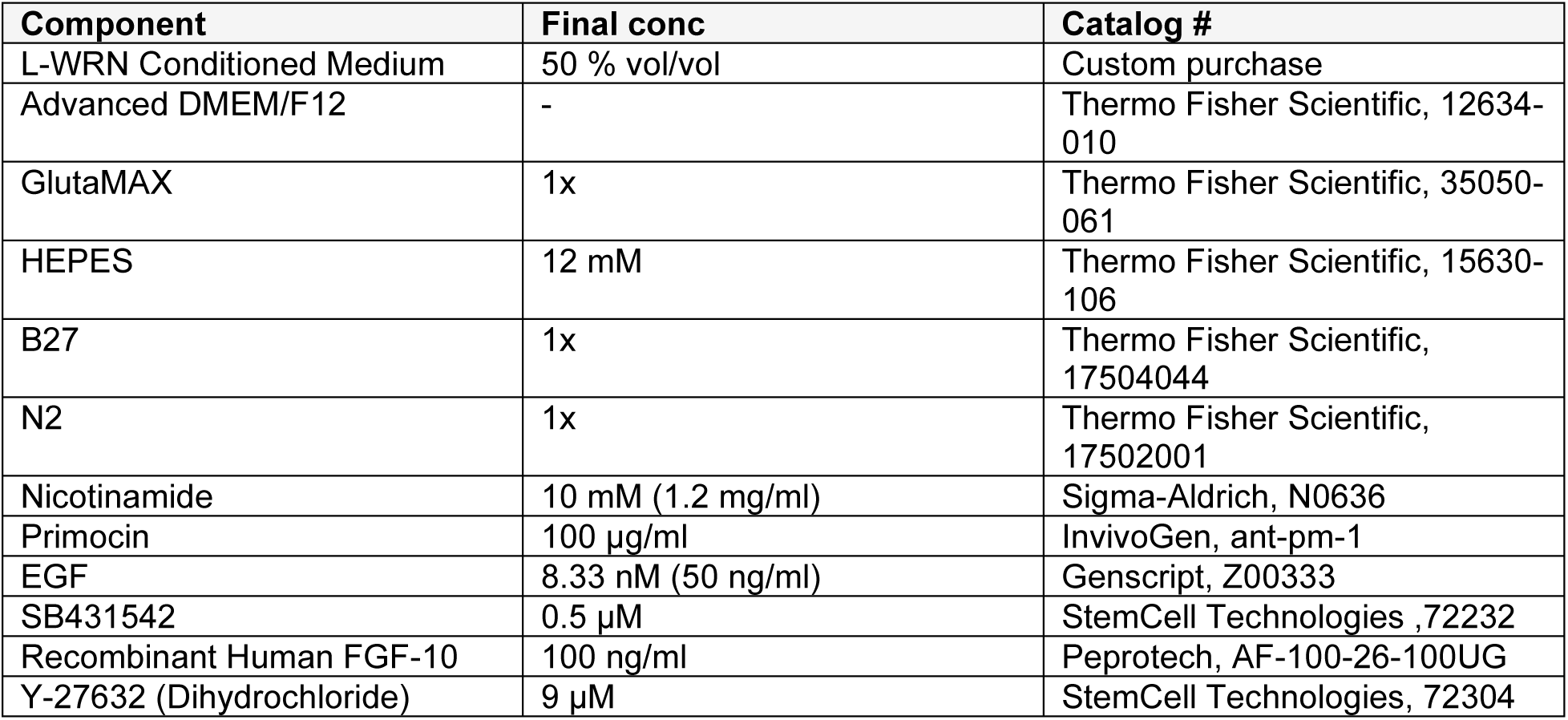
Fallopian Tube WRN-Medium (epithelial)

### Other media formulations

**Supplementary Table S4:** Low serum fibroblast medium composition (note: used in basal chip expansion media)

| Component | Volume/Concentration | Catalog # |
| --- | --- | --- |
| DMEM, high glucose, GlutaMAX | 500 ml | ThermoFisher Scientific, 10-566-016 |
| L-glutamine | 7.5 mM | (ATCC, PCS-201-041 kit component) |
| rh FGF basic | 5 ng/mL | (ATCC, PCS-201-041 kit component) |
| rh Insulin | 5 µg/mL | (ATCC, PCS-201-041 kit component) |
| Hydrocortisone | 1 µg/mL | (ATCC, PCS-201-041 kit component) |
| Ascorbic acid | 50 µg/mL | (ATCC, PCS-201-041 kit component) |
| Fetal bovine serum | 2% | (ATCC, PCS-201-041 kit component) |

### Expansion on chip

**Supplementary Table S5:** Apical Expansion Medium on Chip.

| Component | Content | Catalog # |
| --- | --- | --- |
| Fallopian Tube WRN-Medium | 50% | (see table S3) |
| DermaCult™ Keratinocyte Expansion Medium (with Hydrocortisone) | 50% | StemCell Technologies, 100-0500<br>StemCell Technologies, 07925 |

**Supplementary Table S6:**
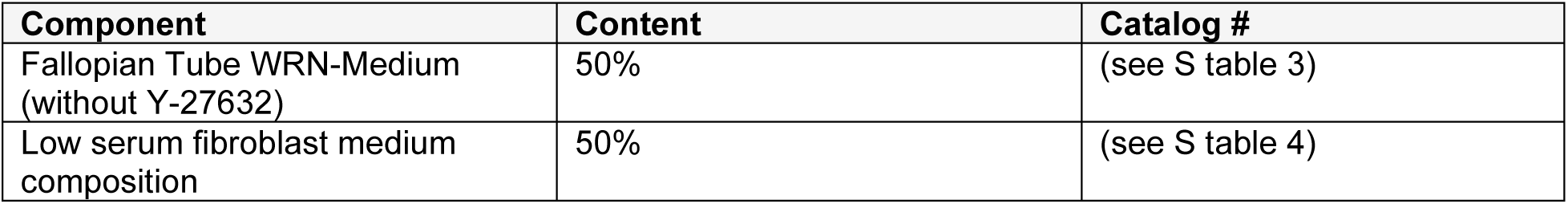
Basal Expansion Medium on Chip.

| Component | Content | Catalog # |
| --- | --- | --- |
| Fallopian Tube WRN-Medium (without Y-27632) | 50% | (see S table 3) |
| Low serum fibroblast medium composition | 50% | (see S table 4) |

### Differentiation on chip

**Supplementary Table S7:** Apical Differentiation Medium on Chip.

| Component | Content | Catalog # |
| --- | --- | --- |
| Sperm wash medium (modified HTF Medium with Human Serum Albumin (5 mg/ml)) | 100% | FUJIFILM Irvine Scientific, NC0231898 |

**Supplementary Table S8:** Basal Differentiation Medium on Chip.

| Component | Content | Catalog # |
| --- | --- | --- |
| GyneCult™ Fallopian Tube Organoid Medium | 100% | StemCell Technologies, 100-1244 |
| Hormones | No hormones (NH)<br>E2<br>E2 + P4 |  |

**Supplementary Table S9:** Sperm Capacitating Medium (c-HTF) Composition.

| Component | Volume/Concentration | Catalog # |
| --- | --- | --- |
| Sperm Washing Medium |  | (FUJIFILM SKU:9983-100ML) |
| Sodium Bicarbonate | 25mM | Gibco™ Sodium Bicarbonate 7.5% solution, 25-080-094 |
| Antibiotics (Primocin™) | 100µg/ml | InvivoGen, ant-pm-1 |

## Supplementary Figures

**Supplementary Figure S1.**
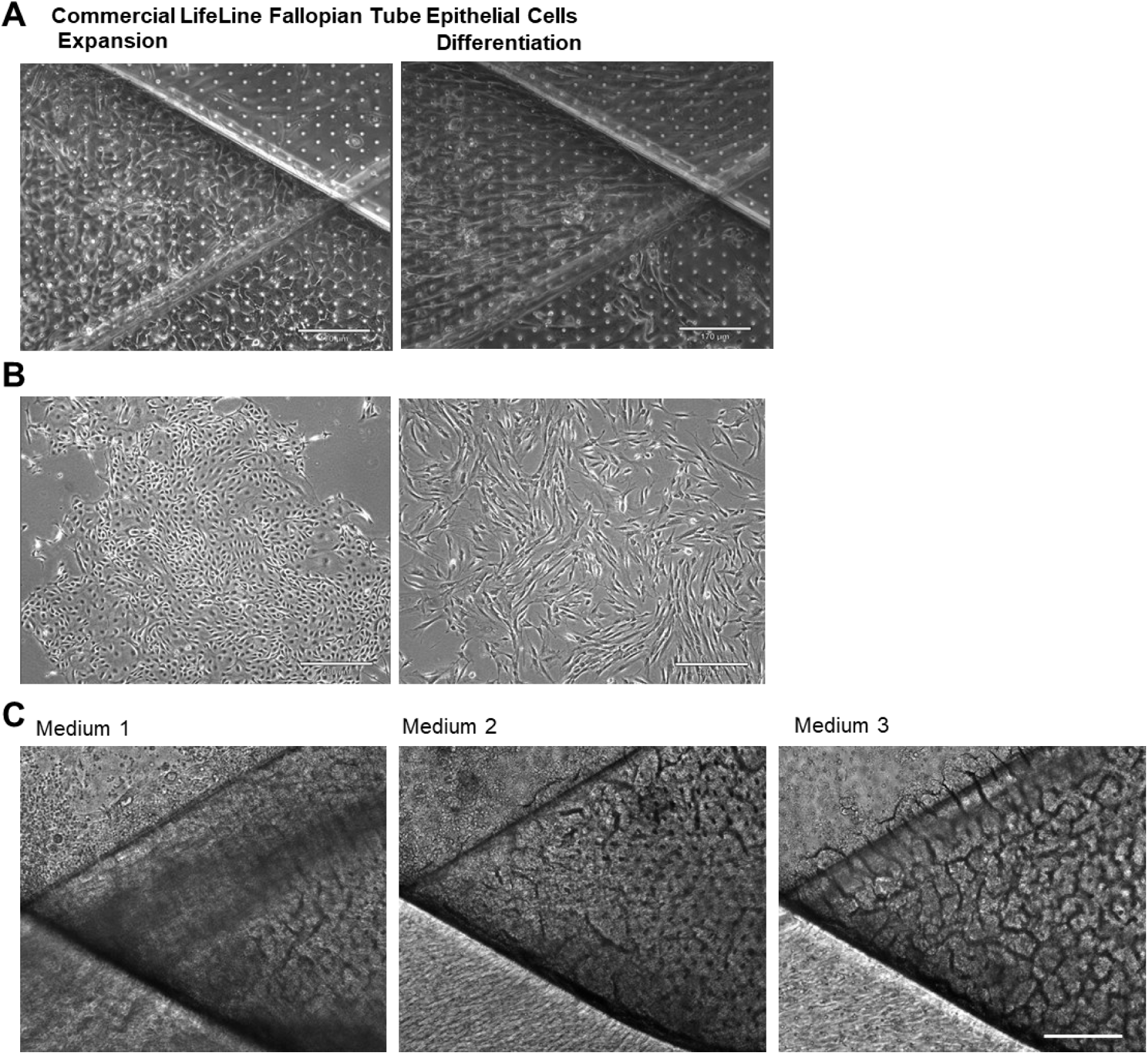
Screening of Cell Sources and Cell Culture Media for Optimal Fallopian Tube Chip Culture. A. Microscopic images showing FT Chip Phenotype using commercially available fallopian tube epithelial cells during the expansion and differentiation stages. (Scale bar, 170 µm). B. Microscopic images depicting ampulla-derived epithelial cells (left) and stromal cells (right) at P0. (Scale bar, 430 µm). C. Microscopic images of FT Chips on day 16 differentiated using three different basal differentiation media, Medium 1(Zhu et al, Biomolecules^1^), Medium 2 (Stejskalova et al, BiorXiv^2^) and Medium 3 (Gynecult^TM^). (Scale bar, 180 µm).

**Supplementary Figure S2.**
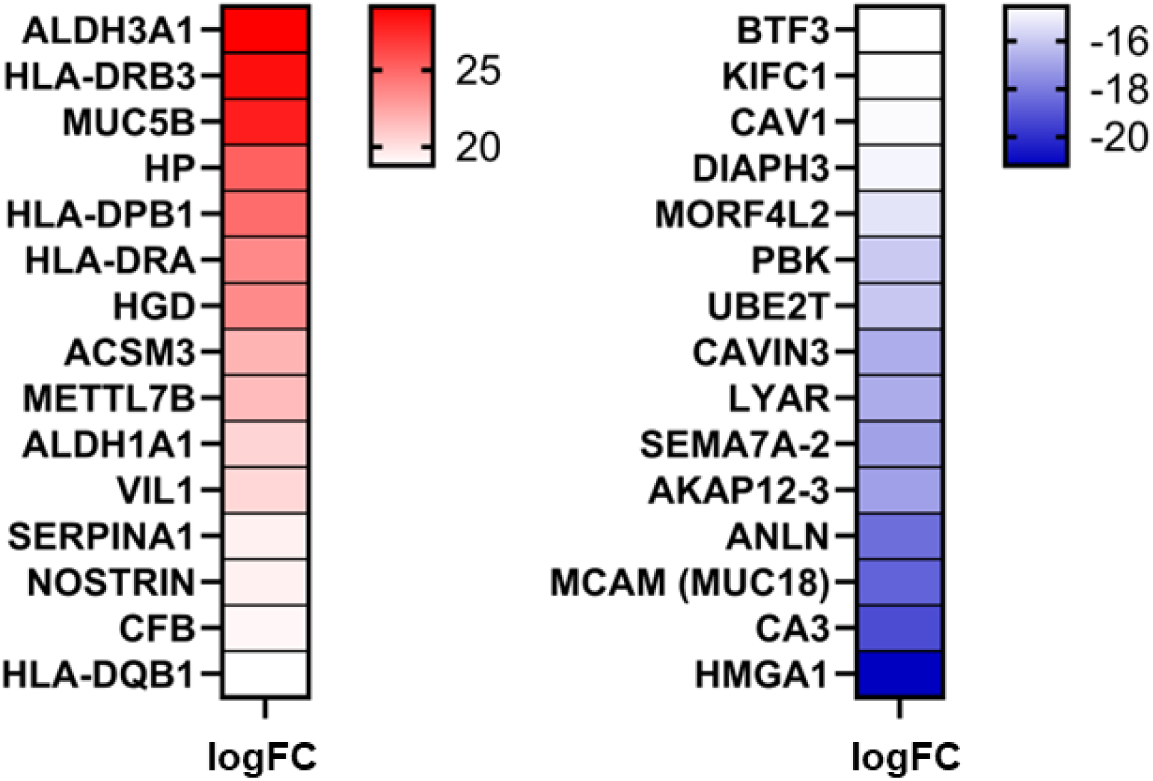
Proteomic comparison of Fallopian Tube Epithelial cells in 2D and matured on Chip (day 16). Left heatmap depicts top 15 most upregulated proteins in the Chip Culture and the right heatmap shows the 15 most differentially downregulated proteins in 2D Culture determine using differential protein expression analysis. logFC represents log fold change. N=one biological replicate, N= 3 technical replicates

**Supplementary Figure S3.**
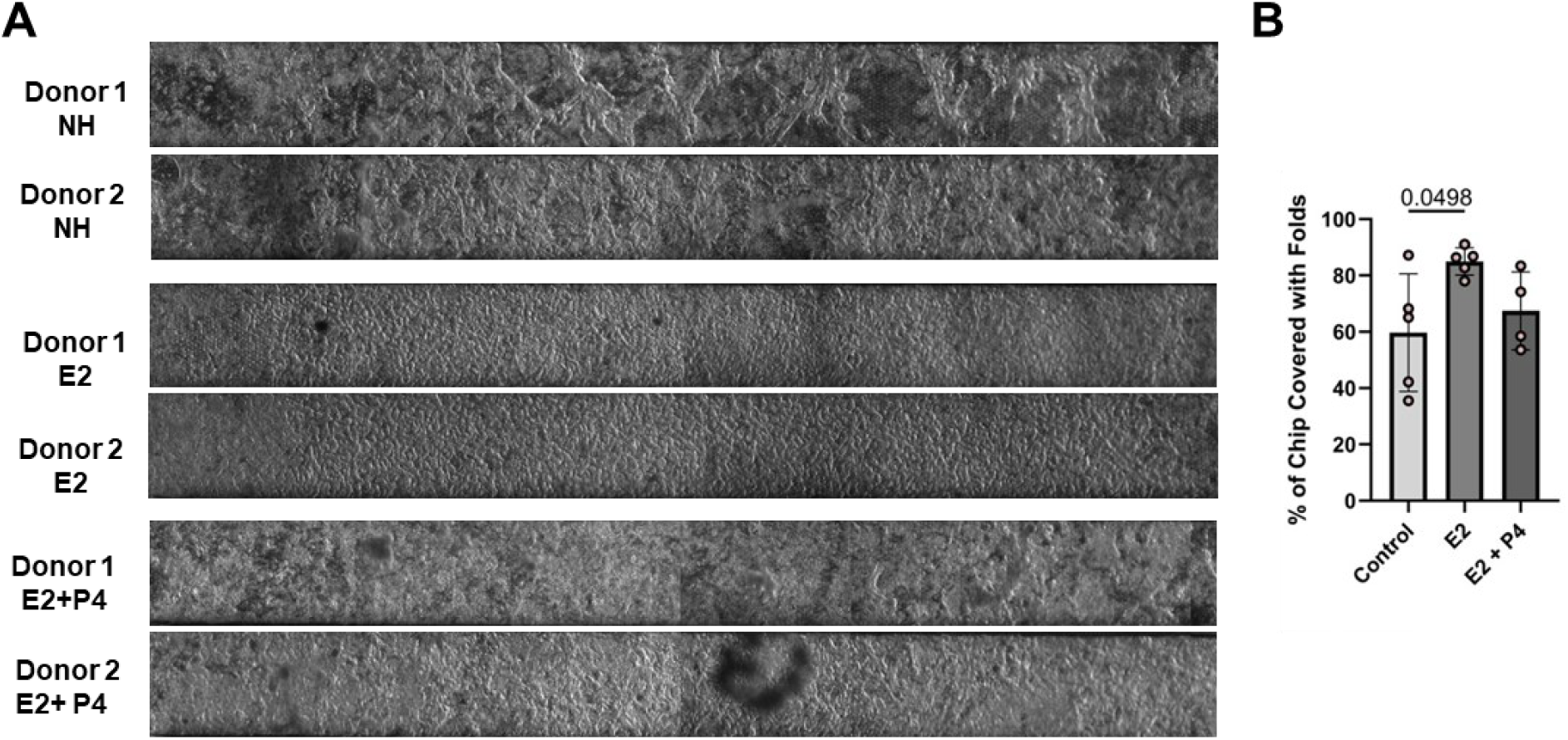
Fold on FT Chips differentiated under different hormonal conditions. **A**. Representative tiled images of the whole FT chips differentiated without added hormones (NH), with 2nM E2 and with 0.1nME2 + 1µM P4. B. Bar Graph depicting the area of chips covered with folds and not flat epithelial monolayer. N=two biological donors, N=4-5 chips. One-way ANOVA was used to determine significance.

**Supplementary Figure S4.**
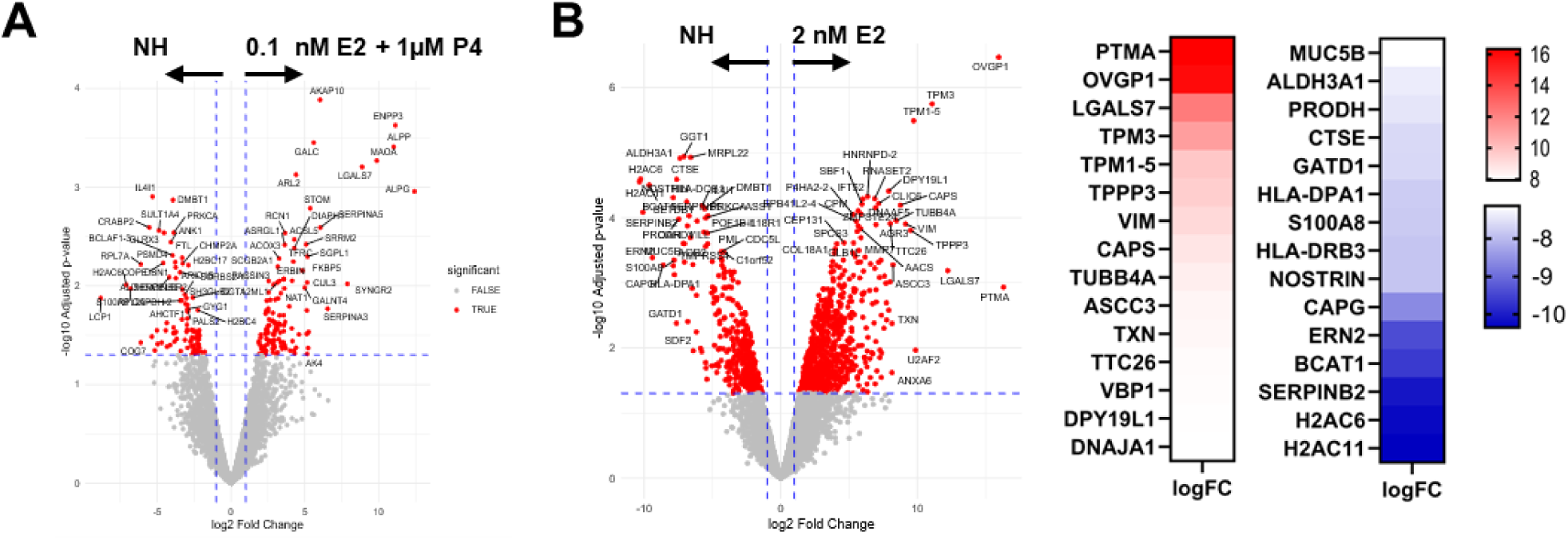
Proteomic comparison of Fallopian Tube Chip epithelia under distinct hormonal treatments. **A.** Volcano plot of differentially expressed proteins between FT Chips perfused basally with 0.1 nME2 + 1µM P4 or differentiation medium alone (NH). **B**. Volcano plot showing differentially expressed proteins between the E2 2nM group and NH groups with heatmap in the middle showing 15 most differentially upregulated proteins in the E2 group and blue heatmap on the right showing 15 most differentially downregulated proteins.logFC signifies log fold change.

**Supplementary Figure S5.**
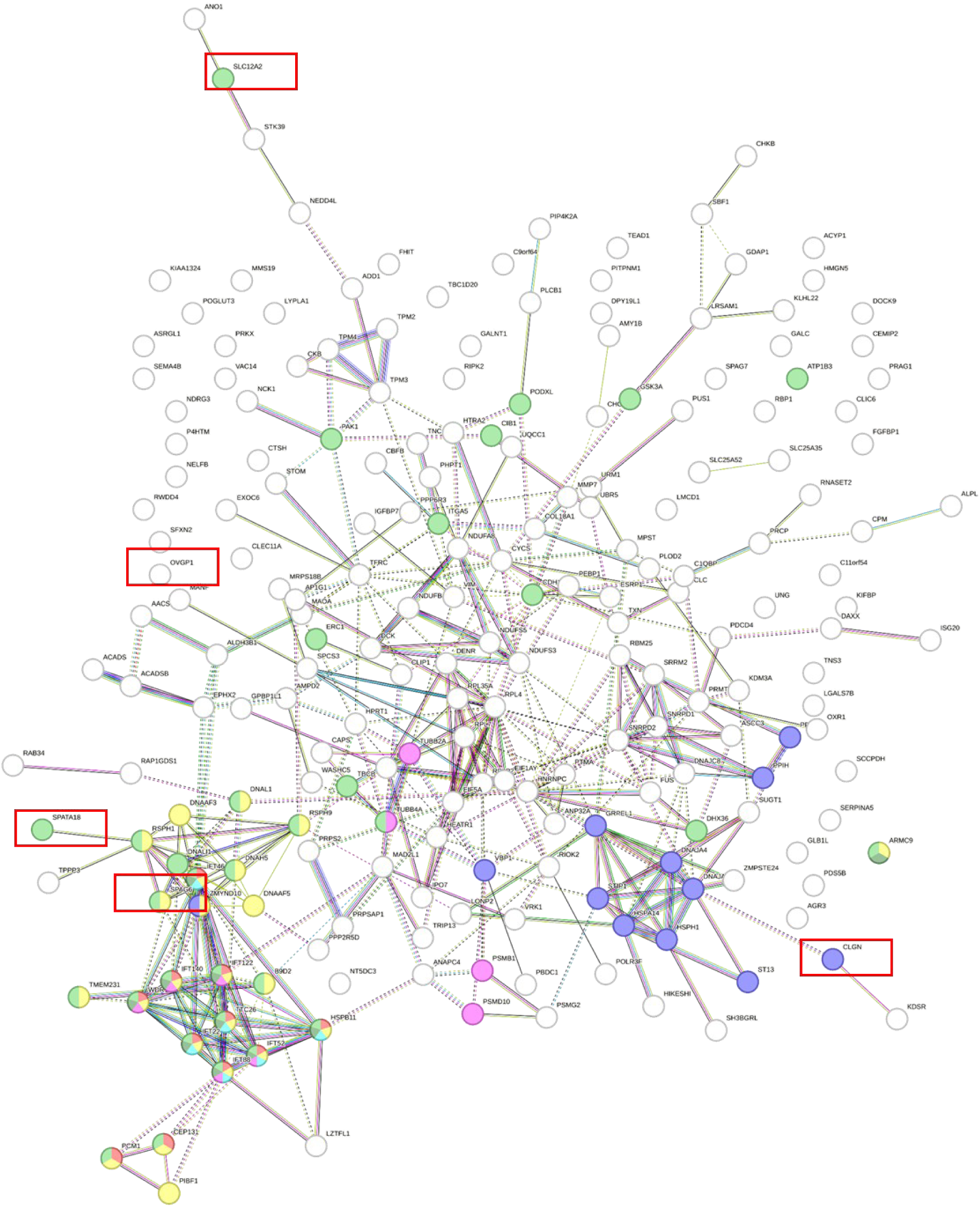
Figure depicting a Protein-protein interaction (PPI) network for proteins that were upregulated in the FT Chip E2 compared to FT Chip NH group generated using StringDB. Red rectangles highlight proteins that have a function in reproduction.

**Supplementary Figure S6.**
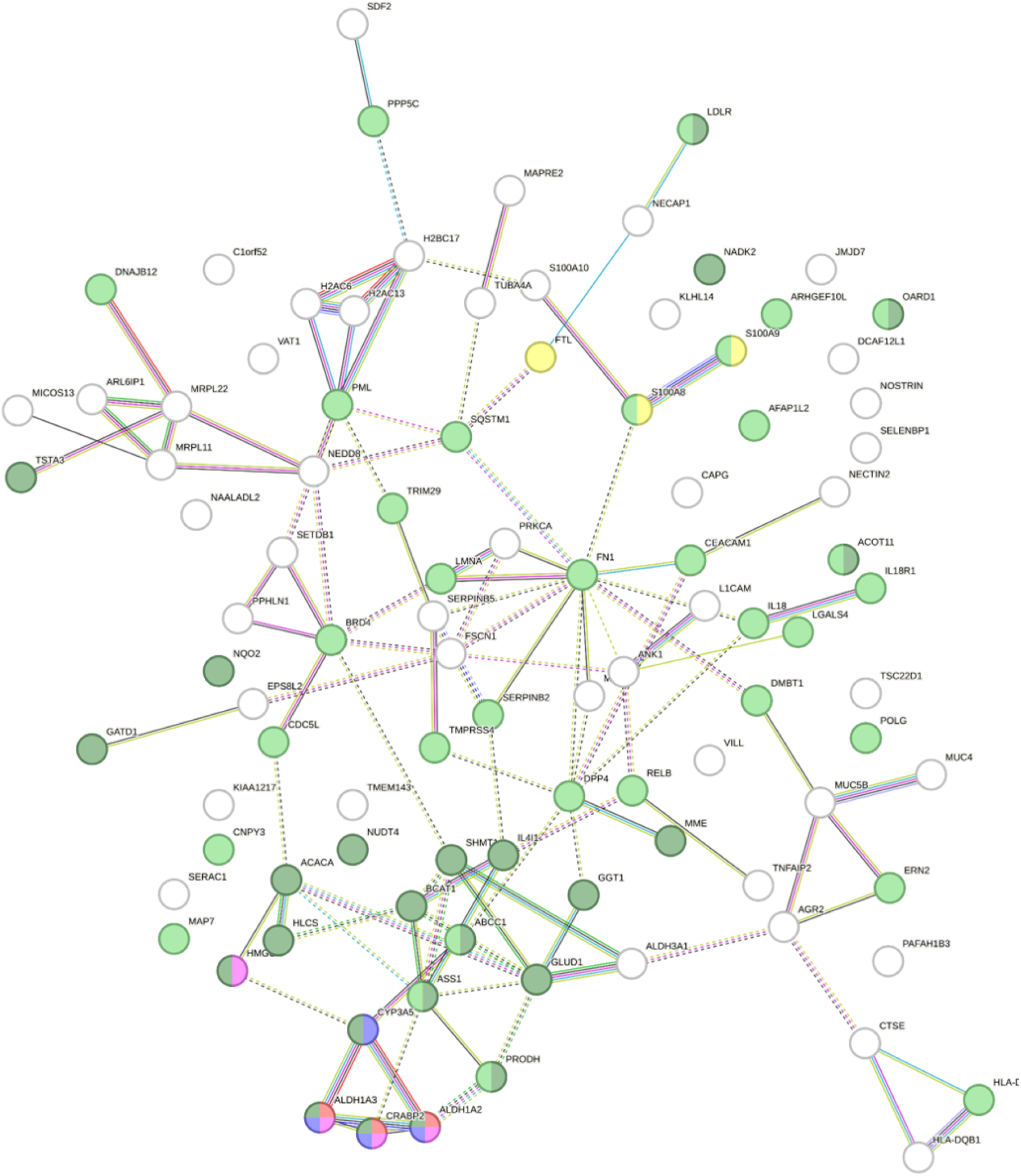
Figure depicting a Protein-protein interaction (PPI) network for proteins that were upregulated in the FT Chip NH compared to FT Chip E2 group generated using StringDB.

**Supplementary Figure S7.**
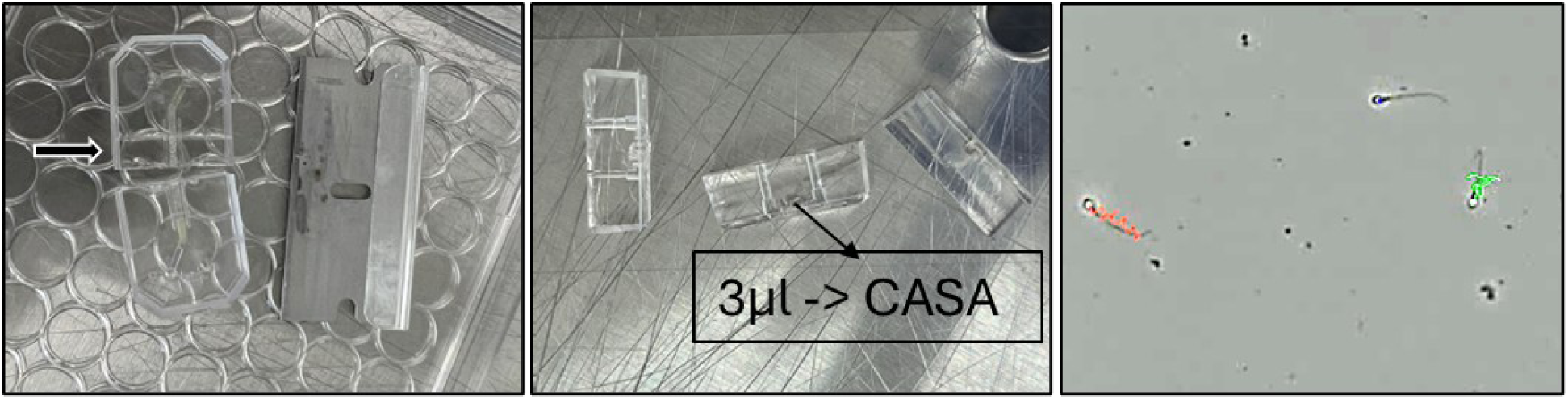
Schematic figure of live-spermatozoa retrieval from the FT Chips. (S1 Chips Emulate). The individual Chips were sliced at the central part of the chip using a razor blade (left). The volume of the apical channel of individual slices placed on glass slides was collected (center) and placed directly on CASA slides (right) and subjected to objective CASA spermatozoa movement analysis. Red tracks correspond to Rapid progressive movement (Type A), green tracks corresponds to Medium progressive movement (type B) and blue tracks correspond to Non-Progressive movement (type C).

